# Nonlytic cellular release of hepatitis A virus requires dual capsid recruitment of the ESCRT-associated Bro1 domain proteins HD-PTP and ALIX

**DOI:** 10.1101/2022.04.26.489493

**Authors:** Takayoshi Shirasaki, Hui Feng, Helen M. E. Duyvesteyn, William G. Fusco, Kevin L. McKnight, Ling Xie, Mark Boyce, Sathish Kumar, Rina Barouch-Bentov, Olga González-López, Ryan McNamara, Li Wang, Adriana Hertel-Wulff, Xian Chen, Shirit Einav, Joseph A. Duncan, Maryna Kapustina, Elizabeth E. Fry, David I. Stuart, Stanley M. Lemon

**Affiliations:** Lineberger Comprehensive Cancer Center, The University of North Carolina at Chapel Hill, Chapel Hill, NC 27599, USA; Division of Structural Biology, The Wellcome Trust Centre for Human Genetics, University of Oxford, Oxford OX3 7BN, UK; Diamond Light Source, Didcot OX11 0DE, UK; Department of Medicine, The University of North Carolina at Chapel Hill, Chapel Hill, NC 27599, USA; Department of Biochemistry and Biophysics, The University of North Carolina at Chapel Hill, Chapel Hill, NC 27599-7292, USA; Division of Infectious Diseases and Geographic Medicine, Department of Medicine, Stanford University School of Medicine, Stanford, California, USA; Department of Microbiology and Immunology, Stanford University School of Medicine, Stanford, California, USA; Chan-Zuckerberg BioHub, San Francisco, CA 94158; Department of Pharmacology, The University of North Carolina at Chapel Hill; Department of Cell Biology & Physiology, The University of North Carolina at Chapel Hill, Chapel Hill, NC 27599, USA; Department of Microbiology & Immunology, The University of North Carolina at Chapel Hill, Chapel Hill, NC 27599, USA

## Abstract

Although picornaviruses are conventionally considered ‘nonenveloped’, members of multiple picornaviral genera are released nonlytically from infected cells in extracellular vesicles. The mechanisms underlying this process are poorly understood. Here, we describe interactions of the hepatitis A virus (HAV) capsid with components of host endosomal sorting complexes required for transport (ESCRT) that play an essential role in release. We show release of quasi-enveloped virus (eHAV) in exosome-like vesicles requires a conserved export signal located within the 8 kDa C-terminal VP1 pX extension that functions in a manner analogous to late domains of canonical enveloped viruses. Fusing pX to a self-assembling engineered protein nanocage (EPN-pX) resulted in its ESCRT-dependent release in extracellular vesicles. Mutational analysis identified a 24 amino acid peptide sequence located within the center of pX that was both necessary and sufficient for nanocage release. Deleting a YxxL motif within this sequence ablated eHAV release, resulting in virus accumulating intracellularly. The pX export signal is conserved in non-human hepatoviruses from a wide range of mammalian species, and functional in pX sequences from bat hepatoviruses when fused to the nanocage protein, suggesting these viruses are released as quasi-enveloped virions. Quantitative proteomics identified multiple ESCRT-related proteins associating with EPN-pX, including ALG2-interacting protein X (ALIX), and its paralog, tyrosine-protein phosphatase non-receptor type 23 (HD-PTP), a second Bro1 domain protein linked to sorting of ubiquitylated cargo into multivesicular endosomes. RNAi-mediated depletion of either Bro1 domain protein impeded eHAV release. Super-resolution fluorescence microscopy demonstrated colocalization of viral capsids with endogenous ALIX and HD-PTP. Co-immunoprecipitation assays using biotin-tagged peptides and recombinant proteins revealed pX interacts directly through the export signal with N-terminal Bro1 domains of both HD-PTP and ALIX. Our study identifies an exceptionally potent viral export signal mediating extracellular release of virus-sized protein assemblies and shows release requires non-redundant activities of both HD-PTP and ALIX.

**Authors’ Summary:** Mechanisms underlying nonlytic release of canonical nonenveloped viruses from infected cells are poorly understood. We show here that release of hepatitis A virus from cells in exosome-like vesicles requires nonredundant activities of two distinct Bro1-domain proteins associated with host cell machinery (ESCRT) for endosomal sorting, HD-PTP and ALIX. We demonstrate both Bro1 domain proteins are recruited to the viral capsid by the pX segment of the 1D capsid protein, and that they act in a non-redundant manner to mediate virus release. Fusing pX to a self-assembling nanocage protein resulted in ESCRT-dependent release mediated by a short pX peptide sequence conserved in hepatoviruses from bats to humans. Mutations within the pX sequence ablate release and result in noncytolytic virus accumulating intracellularly. Our study identifies an exceptionally potent viral export signal mediating extracellular release of virus-sized protein assemblies and shows nonlytic release of quasi-enveloped virus is an ancient evolutionary trait of hepatoviruses.

## Introduction

Viruses from a variety of virus families that are classically considered ‘non-enveloped’ have been found to be released nonlytically from infected cells in extracellular vesicles (EVs), a phenomenon with potentially profound consequences for pathogenesis and host immune response [1–4]. The size of the EVs containing these quasi-enveloped viruses, the number of viral capsids each contains, and the host proteins with which they are associated vary widely. This likely reflects distinct mechanisms of biogenesis, none of which are well understood. Several pathogenic viruses classified within different genera of the *Picornaviridae*, a large and diverse family of positive-strand RNA viruses [5], are released from cells in this fashion. EVs containing poliovirus and coxsackivirus (genus *Enterovirus*) are large (300-920 nm across) and contain numerous capsids, whereas EVs containing hepatitis A virus (HAV, genus *Hepatovirus*) are smaller (50-110 nm) and contain only 1-3 capsids [1, 2, 6]. Enterovirus and encephalomyocarditis virus (genus *Cardiovirus*) EVs are enriched with the autophagy-related protein LC3 and may be derived from autophagosomes, whereas HAV is released in EVs that resemble exosomes in size and protein composition and likely originate in multivesicular endosomes (MVEs) [1, 2, 4, 7]. Thus, even within a single virus family, it is likely that diverse signals drive the quasi-envelopment of viral capsids.

EVs containing HAV have specific infectivity (infectious particles/genome number) similar to naked, nonenveloped virions, despite the absence of any virus-encoded protein on their surface [1, 7]. These quasi-enveloped virions (eHAV) enter cells via interactions with cell surface phosphatidylserine receptors, and require endolysosomal ganglioside receptors shared with naked, non-enveloped HAV (nHAV) [8, 9]. They are the only form of virus detected in the blood during acute hepatitis A [1]. Canonical nonenveloped picornaviral virions are produced in the biliary track, where membranes are stripped from quasi-enveloped eHAV shed from the liver [10]. The membranes surrounding the capsid in eHAV cloak it from the immune system, and likely contribute to the stealth-like nature of early HAV infection [1, 11]. Replication of HAV is noncytopathic *in vivo* [12] and its nonlytic release in vesicles represents a defining feature of the virus.

Multiple proteins associated with endosomal sorting complexes required for transport (ESCRT) have been identified in extracellular eHAV vesicles [7]. These include ALG2-interacting protein X (ALIX, also known as PDCD6-interacting protein), a Bro1-domain protein that is a key player in MVE formation, cytokinesis, and the budding of conventional enveloped viruses from host cell membranes [13, 14]. Multiple components of ESCRT-III complexes are also present, including CHMP4B (charged multivesicular body protein 4B), CHMP1A, CHMP1B and IST1 (Ist1 homolog 1) [7]. RNAi-mediated depletion of ALIX, IST1, and CHMP2A suppresses eHAV release from infected cells, indicating that ESCRT is required for eHAV biogenesis [1, 7]. The VP2 capsid protein of HAV contains tandem YPX_1/3_L ‘late domains’ similar to linear peptide motifs in structural proteins of enveloped viruses that are known to bind ALIX and recruit ESCRT [15, 16]. Such late domains are essential for the budding of enveloped viruses from host cell membranes [17, 18]. Mutational data support the importance of the VP2 YPX_1/3_L motifs in eHAV release [16], although paradoxically the residues involved present little accessible surface area in an X-ray model of the mature, extracellular capsid [19].

Other data point to the involvement of a poorly understood, 8 kDa C-terminal extension of the HAV VP1 capsid protein known as ‘pX’ in eHAV biogenesis (**Fig. 1A**). Originally identified in virion assembly intermediates and found more recently in extracellular eHAV particles [1, 20], the pX sequence is unique to hepatoviruses. pX is rapidly cleaved from VP1 upon loss of the eHAV membrane and is not present in nHAV [1]. The N-terminal 27 residues of pX regulate folding of upstream capsid sequence and are required for capsid protein pentamer formation early in virion assembly [21, 22]. Less is known about the C-terminal 43 residues of pX, which can sustain large deletions without loss of infectivity [23]. Previous studies suggest pX forms a complex with ALIX, and show that deletion of the C-terminus prevents quasi-envelope virus release [24]. Here, we show pX also interacts with tyrosine-protein phosphatase non-receptor type 23 (HD-PTP), an ALIX paralog associated with sorting of ubiquitylated cargo into MVEs [25]. We demonstrate that pX binds directly to the N-terminal Bro1 domains of both HD-PTP and ALIX, and that these two ESCRT-associated proteins function nonredundantly in a coordinated fashion to mediate the release of quasi-enveloped virus. Our results expand the range of ESCRT-associated proteins involved in virus release, define a potent viral export signal in pX, and show quasi-envelopment to be conserved feature of hepatoviruses recovered from mammalian species ranging from bats to humans.

**Figure 1.**
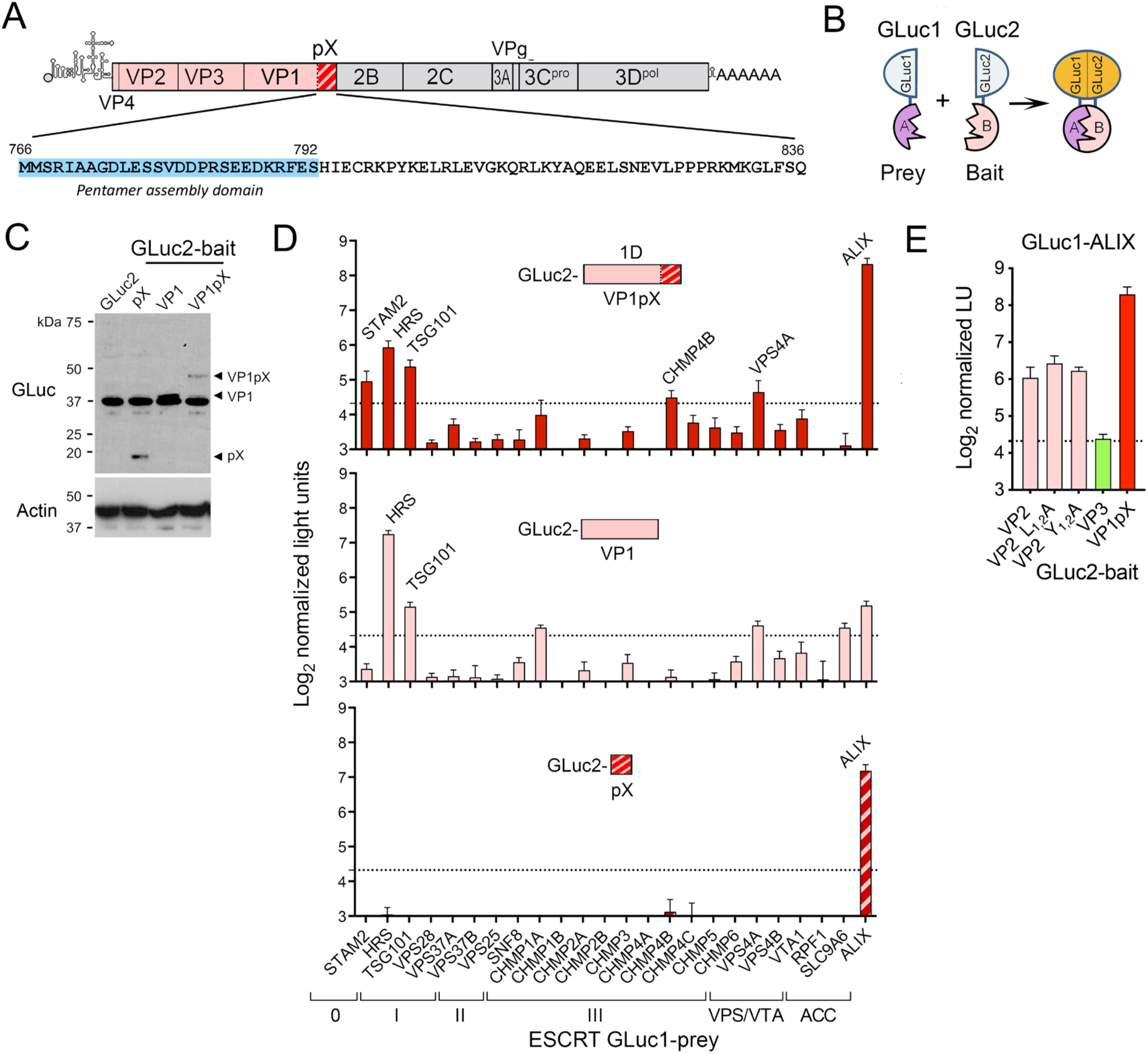
Protein fragment complementation screen for VP1pX interactions with ESCRT. **(A)** HAV genome organization: capsid proteins are in color with the pX hatched. Below are pX sequences of p16 virus and related mutants. **(B)** Protein-fragment complementation assay with ‘bait’ and ‘prey’ proteins fused to N-terminal GLuc1 and C-terminal GLuc2 *Gaussia princeps* luciferase fragments [28]. (**C**) Representative GLuc immunoblot showing expression levels of GLuc2 bait proteins fused to VP1pX (1D), VP1, or pX in 293T cells. Actin included as a loading control. (**D**) Screen for interactions of (*top*) VP1pX, (*middle*) VP1, and (*bottom*) pX with ESCRT components. Normalized light units (LU) >20 (dashed line) exceed values from a large panel of control prey proteins [27]. Acc: ESCRT accessory protein. **(E)** Results of protein-fragment complementation assay assessing interactions of GLuc1-ALIX prey with VP2, VP2 late domain mutants with Ala substitutions of Leu (L_1-2_A) and Tyr residues (Y_1-2_A) (*14*), VP3, or VP1pX fused to GLuc2 as bait. Data in C and E are means from 2 independent experiments, each with 3 technical replicates.

## Results

### A protein-fragment complementation screen reveals VP1pX interactions with ESCRT proteins

To better understand how pX functions in eHAV release, we screened the entire 1D capsid protein (VP1pX) for interactions with ESCRT-associated proteins using a sensitive *Gaussia princeps* luciferase (GLuc) protein-fragment complementation assay that is capable of detecting weak and/or transient protein-protein interactions [26–28]. Unprocessed VP1pX protein, fused as ‘bait’ to the C-terminal half of GLuc (‘GLuc2’), was transiently co-expressed in 293T cells with a panel of 24 ESCRT-associated proteins fused as ‘prey’ to the N-terminus of GLuc (‘GLuc1’) (**Fig. 1B,C**). A robust interaction signal was generated with ALIX, and to a lesser extent with proteins associated with ESCRT-0 (HRS and STAM2) and ESCRT-I (TSG101) (**Fig. 1D** ***top***). Subsequent screens using isolated VP1 and pX polypeptides as bait suggested ALIX interacts primarily with pX, whereas the interaction with HRS and TSG101 mapped to VP1 (**Fig. 1D** ***bottom, middle***). Additional studies with GLuc2-VP2 bait were consistent with the previously reported ALIX-interacting YPX_3_L late domains in VP2 [16], although the signal was not reduced by single Ala substitutions of Tyr or Leu residues in each of the domains (**Fig. 1E**). By contrast, VP3 showed no evidence of interacting with ALIX. These data are consistent with ESCRT-recruitment signals residing in VP1pX as well as VP2, with pX interacting either directly or indirectly with ALIX as reported previously [24].

### pX drives extracellular release of computationally-designed nanocage assemblies

HAV replication is slow and inefficient in cell culture, complicating the functional analysis of viral proteins. Thus, to assess the role of pX in eHAV release, we fused the pX sequence to the C-terminus of a 203 amino acid *de novo* designed protein that self-assembles into a 60-copy, 25 nm wire cage dodecahedron [29] (**Fig. 2A**). These engineered protein nanocages (EPN) have been shown to be secreted from cells within vesicles by an ESCRT-dependent process when the protein is fused at its C-terminus to the p6^Gag^ protein of human immunodeficiency virus (HIV-1) that contains late domains interacting with TSG101 and ALIX [30]. Secretion is also dependent upon the EPN protein being linked at its N-terminus to residues 2-6 of Gag that contains a myristoylation signal mediating association with membranes [30]. Fusing pX to the C-terminus of the nanocage protein (‘EPN-pX’) in lieu of p6^Gag^ did not interfere with nanocage assembly, and resulted in nanocages being visibly present within membrane-limited vesicles released from transfected 293T cells (**Fig. 2B**). Detergent treatment was required to immunoprecipitate EPN-pX released by the transfected cells, consistent with its sequestration within vesicles (**Fig. 2C**). Furthermore, abundant EPN-pX could be recovered from beneath a 20% sucrose cushion following centrifugation of extracellular fluids, confirming the particulate nature of the secreted nanocage **(Fig. 2D, lanes 1 and 13)**. A 6.7 Å 3D structure of these nanocages determined by cryo-electron microscopy was very similar to that reported previously [29] **(Fig. S1A-C)**. However, additional density was evident in the vicinity of the C-terminus of the nanocage protein, facing towards the inside of the nanocage and consistent with pX **(Fig. S1D,E)**. Release was suppressed by co-expression of a dominant-negative VPS4A mutant (E228Q) (**Fig. 2D**, compare lanes 1 and 7), confirming ESCRT-dependence [30]. Nanoparticle tracking analysis revealed a small increase in the median size of EVs released by 293T cells expressing EPN-pX (127 nm) compared with control cells (110 nm) **(Fig. S1F,G)**, but little change in overall EV concentration **(Fig. S1H)**. Thus, EPN-pX expression minimally perturbed the background release of exosomes and other microvesicles from the cells.

**Figure 2.**
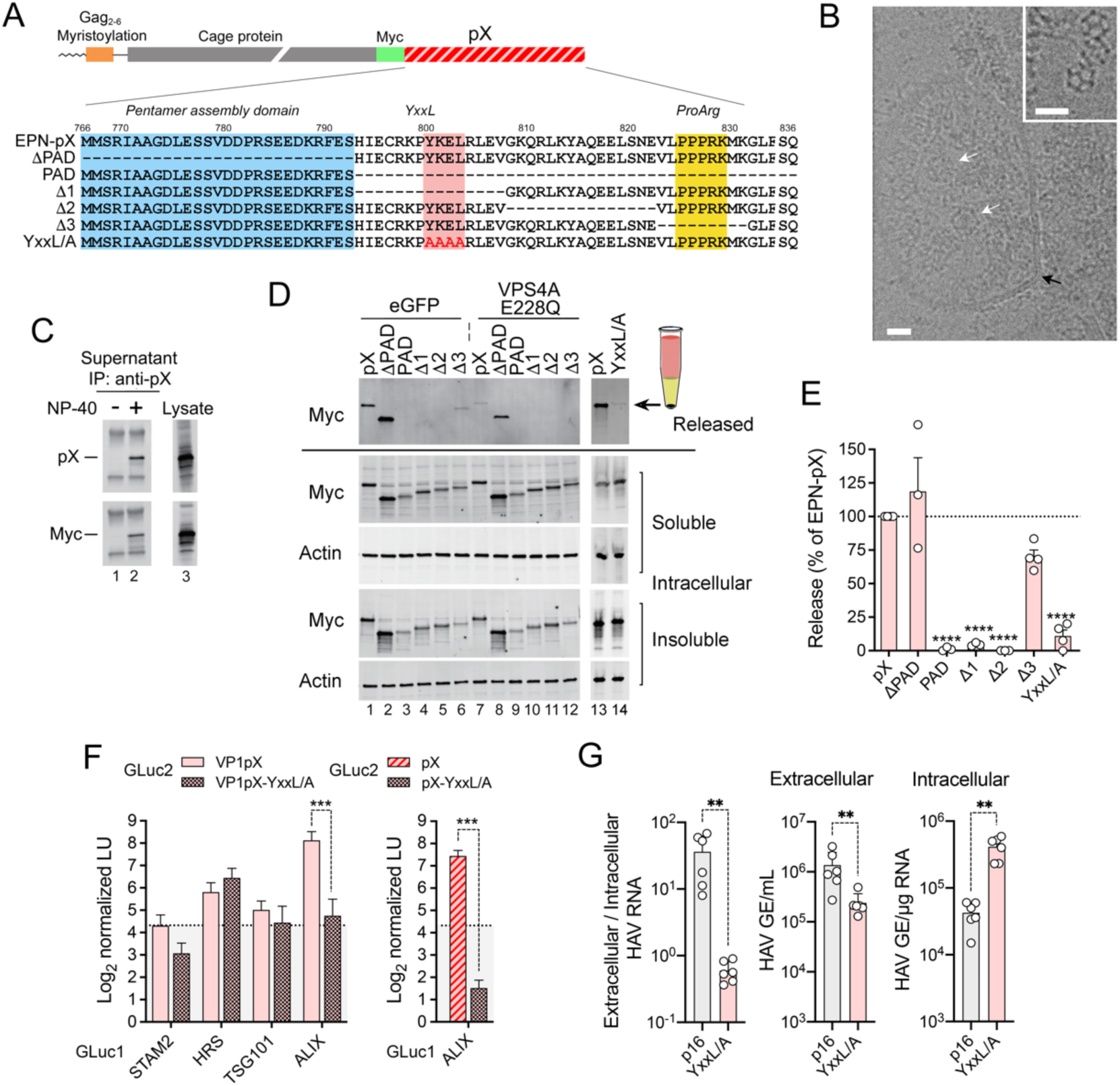
pX mediates ESCRT-dependent extracellular release of assembled EPN-pX nanocages and noncytopathic p16 virus. **(A) (***top*) EPN-pX, in which pX is fused to the C-terminus of the nanocage protein with an intervening Myc tag. (*bottom*) pX sequence in EPN-pX and related mutants. The pentamer assembly domain (PAD) [21] and YxxL motif are indicated. Δ1, Δ2 and Δ3 deletions were described previously [23]. **(B)** Representative electron micrograph showing multiple EPN-pX nanocages (*white arrows*) within a membrane-bound vesicle (*black arrow)* in extracellular fluids of transfected 293T cells. Scale bar = 20 nm. The inset shows nanocages in extracellular fluids treated with CHAPS. Scale bar = 30 nm. (**C**) pX and Myc immunoblots of anti-pX immunoprecipitates of extracellular fluids, with and without NP-40 treatment, versus lysate from EPN-pX transfected cells. **(D)** Myc immunoblot of released EPN protein recovered following centrifugation through a 20% sucrose cushion, and soluble and insoluble intracellular EPN proteins expressed by cells transfected with EPN-pX and related constructs. Cells were co-transfected with vectors expressing eGFP (control, lanes 1-6) or the E228Q VPS4A mutant (lanes 7-12). (**E**) Quantitation of nanocage protein release normalized to intracellular protein expression. Results are means ±S.E.M. from 3 independent experiments. \*\*\*\**p*<0.0001 by ANOVA with Sidak’s multiple comparison test. **(F)** Protein-fragment complementation assay with (*left*) VP1pX and (*right*) pX GLuc2 baits with and without YxxL/A substitutions and GLuc1 prey fused to ESCRT components. Results are means ± S.D. from two independent experiments, each with 3 technical replicates. ***p<0.001 by two-way ANOVA with Sidak’s multiple comparison test. **(G)** Extracellular and intracellular HAV RNA, and extracellular/intracellular RNA ratios, 12 days after transfection of cells with p16 or p16-YxxL/A RNA. Data are from n=6 cultures from two transfections, and are representative of multiple independent experiments. **p<0.01 calculated by Mann-Whitney test.

EPN-pX was released from cells as efficiently as EPN fused to p6^Gag^ **(Fig. S2A).** Since EPN-p6^Gag^ release requires both TSG101-interacting ‘PTAP’ and ALIX-interacting ‘YPX_3_L’ late domains of p6^Gag^ [30] **(Fig. S2A)**, this indicates that pX is capable of functionally replacing both p6^Gag^ late domains. By contrast, EPN was not released when fused to HAV VP2 sequence containing tandem YPX_3_L late domains that interact only with ALIX [16] **(Fig. S2B)**.

### EPN-pX release requires an association with intracellular membranes

In addition to self-assembly and ESCRT recruitment, the release of EPN-p6^Gag^ requires a myristoylation signal at its N-terminus that drives its association with membranes [30]. EPN-pX release was similarly dependent on this myristoylation signal **(Fig. S2C)**, indicating that pX does not by itself mediate an association with intracellular membranes. The VP4 capsid protein of some picornaviruses is N-terminally myristoylated and plays an important role in membrane penetration during cell entry [31]. By contrast, HAV VP4 is small (only 21-23 amino acids) and lacks a myristoylation signal, yet associates with membranes through its N-terminus [32]. EPN-pX constructs in which the N-terminal 9 or 11 residues of VP4 replaced the Gag myristoylation signal of HIV-1 were actively released, albeit not as efficiently as EPN-pX with the Gag signal **(Fig. S2D)**. Treating cells with IMP-1088, an inhibitor of N-myristoyltransferases (NMT1 and NMT2) [33], reduced the release of EPN-pX containing the N-terminal Gag signal, but had little impact on EPN-pX constructs containing N-terminal VP4 sequence **(Fig. S2D** lanes 6-10, *inset*). These results support previous studies indicating VP4 associates with membranes independently of myristoylation [32], and suggest VP4 could provide the membrane association required for ESCRT-mediated capsid export. However, VP4 is located internally within the mature, extracellular HAV capsid [34]. Dynamic ‘breathing’ is known to occur in the capsids of other picornaviruses [35], but it is uncertain whether this would be sufficient to externalize VP4 and allow it to drive an association of the HAV capsid with membranes.

### A YxxL motif in pX is crucial for nanocage export and eHAV release

Deleting the N-terminal 27 amino acid pentamer assembly domain of pX [21] (‘EPN-ΔPAD’) had neglible impact on EPN-pX release, whereas deleting the C-terminal 44 amino acids, leaving only the pentamer assembly domain, completely abolished release (‘EPN-PAD’, **Figs. 2A,D,E**). This localizes the responsible export signal to the C-terminal half of pX. Three deletions within this sequence that were shown previously to limit HAV spread in cell culture [23] had variable effects on EPN-pX release. Two 15-amino acid deletions (‘Δ1’ and ‘Δ2’) completely ablated release, whereas a 9-amino acid deletion (‘Δ3’) near the C-terminus of pX reduced secretion by 30 ±9.8 % (**Figs. 2A,D,E**). The sequence deleted in the Δ1 mutant contains a YxxL motif (YKEL beginning with Tyr^800^) resembling late domains found in arenavirus and paramyxovirus proteins that facilitate budding of these enveloped viruses [36–38] (**Fig. 2A**). Replacing the residues in this motif with 4 serial Ala substitutions eliminated EPN-pX release (‘YxxL/A’ mutant, **Figs. 2A,D,E**), and also reduced interactions of Gluc2-VP1pX and GLuc2-pX bait with GLuc1-ALIX prey in the protein fragment complementation assay (**Fig. 2F**). Consistent with these results, a mutant virus with similar Ala substitutions in the YxxL motif (‘p16-YxxL/A’ virus) replicated well, but was severely impaired in egress resulting in its accumulation within infected cells (**Fig. 2G**). Intracellular viral RNA copy numbers were increased approximately 10-fold, whereas the ratio of extracellular to intracellular viral RNA was reduced by approximately 60-fold. Taken collectively, these data support the existence of a potent export signal centered within the pX sequence that is essential for release of quasi-enveloped virus.

### The pX export signal is functionally conserved in bat hepatoviruses

To gain a fuller understanding of the pX sequence required for viral release, we compared the pX sequence in human HAV (*Hepatovirus A*) with pX sequences in 8 other non-primate hepatovirus species [39]. Although pX is highly conserved among *Hepatovirus A* strains from humans and nonhuman primates, it is very divergent in hepatoviruses identified in seals, rodents, bats and other small mammals [40, 41] (**Figs. S3, S4A**). Nonetheless, a constraint-based sequence alignment revealed a YxxL motif in 21 of 24 hepatovirus sequences recovered from 18 different mammalian host species **(Fig. S3)**. In 2 other viruses, the Leu residue in the motif was replaced with similarly hydrophobic Val or Ala residues. Multiple residues downstream of the YxxL motif were also highly conserved in sequences from 22 of these hepatoviruses, revealing a broadly conserved pX segment extending between Tyr800 of the YxxL motif and Leu819 in human HAV **(Fig. S3)**. This conserved pX segment spans the Δ1 and Δ2 deletions that ablated EPN-pX release (**Fig. 2A,D**), and exists in 8 of 9 hepatovirus species (all but *Hepatovirus G*). Infectious hepatoviruses have yet to be isolated in cell culture from species other than *Hepatovirus A*, but the sequence conservation evident within this segment of pX suggests it has a similar role in capsid quasi-envelopment in other hepatovirus species.

To test this hypothesis, we replaced the pX sequence in EPN-pX with the pX sequence of M32 virus (*Hepatovirus H*), which was recovered from the African straw-colored fruit bat *Eidolon helvum*, and SMG18520 virus (‘SMG’, *Hepatovirus C*), recovered from the long-fingered bat *Miniopterus manavi* in Madagascar [40]. These bat pX sequences share only 39-42% amino acid identity (56-58% similarity) with pX of *Hepatovirus A* **(Figs. 3A, S4A)**. Despite lower expression of soluble proteins, both EPN-M32pX and EPN-SMGpX were released from cells with 50-80% of the efficiency of EPN-pX (calculated from immunoblot intensities of released versus soluble intracellular protein) (**Fig. 3B**). As with EPN-pX, EPN-M32pX release was reduced or eliminated by co-expression of the dominant negative VPS4A-E228Q mutant, indicating that release was ESCRT-dependent **(Fig. S4B)**. ALIX was also increased in abundance in extracellular, particulate EPN-M32pX and EPN-pX fractions recovered from beneath a sucrose cushion **(Fig. S4B, lanes 2,3)**, indicating that ALIX is recruited to these nanocage assemblies during export. Taken collectively, these data indicate that pX-directed nonlytic virus release is an ancient and functionally conserved attribute of hepatoviruses recovered from mammalian hosts separated by millions of years of evolution.

**Figure 3.**
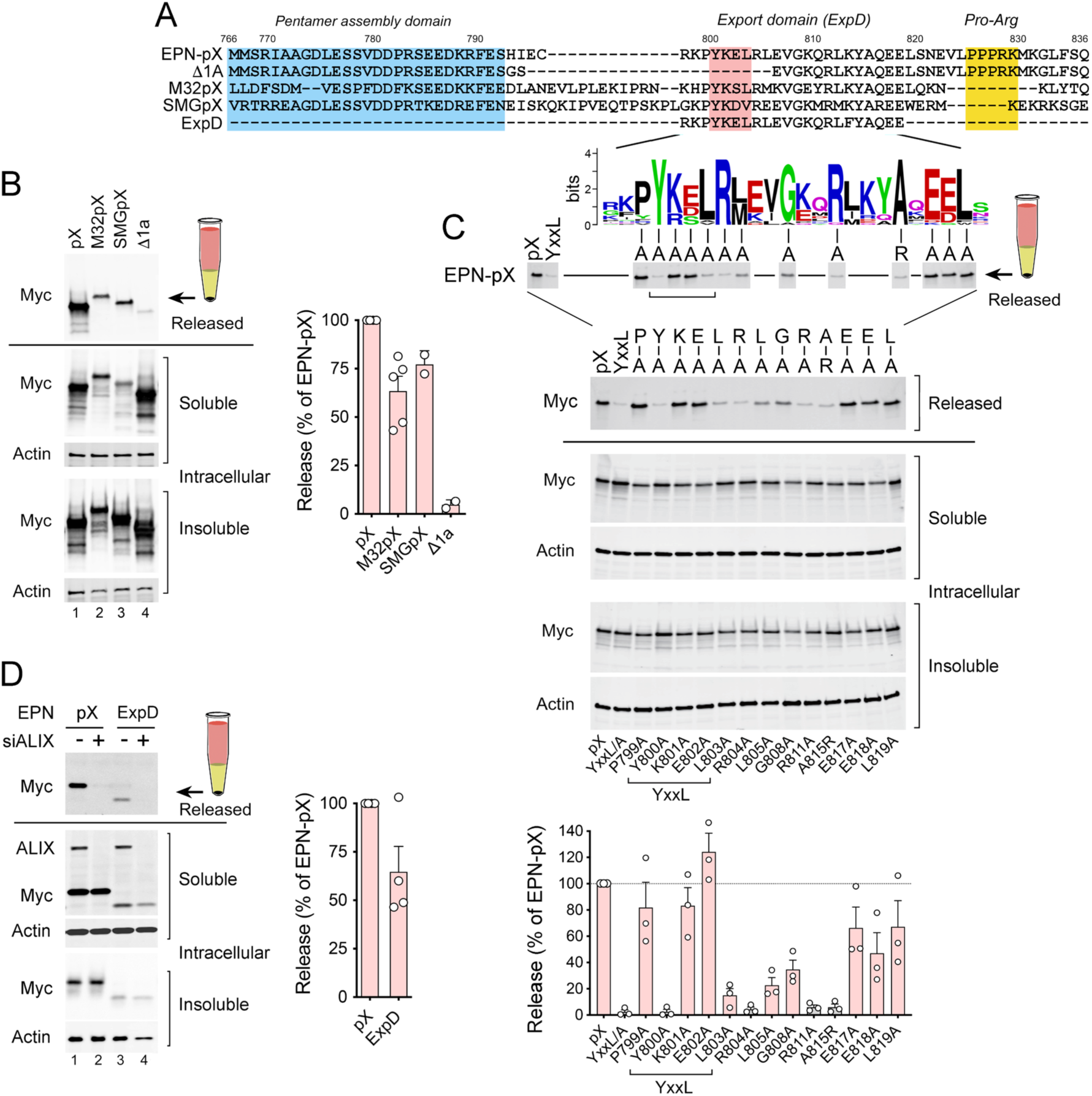
A conserved centrally-located pX peptide sequence, ExpD, mediates ALIX-dependent nanocage release. (**A**) Wild-type and mutant pX sequences of human p16 virus aligned with bat virus pX sequences of M32 (*Hepatovirus H*), recovered from *Eidolon helvum,* and SMG18520 (‘SMG’, *Hepatovirus C*), recovered from *Miniopterus manavi* [40]. The N-terminal pentamer assembly domain, central ExpD sequence, and C-terminal Pro-Arg motif are shaded. Below is shown a WebLogo for ExpD derived from an alignment of 22 pX sequences from different mammalian hepatoviruses (see **Fig. S3**). (**B**) Immunoblots showing released EPN-pX, EPN-M32pX, and EPN-SMGpX nanocages recovered from beneath a sucrose cushion, and related soluble and insoluble intracellular EPN proteins. EPN-Δ1a was included as a negative control. Quantitation is shown on the right, with mean p16 and bat pX nanocage release normalized to intracellular protein expression, ± S.E.M., n=2-5. **(C)** Alanine scanning of conserved amino acid residues within EPN-pX using the nanocage release assay. Shown below are immunoblots of released and intracellular EPN proteins, and at the bottom quantitation of immunoblots in 3 independent experiments. Brackets indicate residues within the YxxL motif. (**D**) Immunoblots showing released and soluble/insoluble intracellular EPN-pX and EPN-ExpD proteins following transfection with ALIX (*PDCD6IP*)-targeting (+) or non-targeting (-) scrambled siRNAs. Immunoblotting of soluble intracellular proteins was with anti-Myc and anti-ALIX antibodies. Quantitation of EPN-ExpD release is shown on the right.

To better define the export signal in pX, we created single Ala substitutions at residues in EPN-pX that are conserved in both the human and bat viruses. Substitutions at Tyr800, Leu803, Arg804, Leu805, Gly808, and Arg811 (*Hepatovirus A* numbering) substantially ablated nanocage release, as did substituting Arg for the conserved Ala815 (**Fig. 3C**). Fusing the conserved sequence at the center of pX (residues 797-820) to EPN (‘EPN-ExpD’) resulted in release of the nanocage with 64 ±26% s.d. the efficiency of EPN-pX (**Fig. 3D**), which is comparable to EPN-Δ3 (**Fig. 2E**). Like EPN-pX, EPN-ExpD release was reduced by siRNA-mediated depletion of ALIX (**Fig. 3D**). These results thus define a conserved ESCRT-interacting export signal comprised of YxxLR[L/M]xxGxxRxxxA at the center of pX that is both necessary and sufficient for ALIX-dependent nanocage release.

### Label-free quantitative proteomic analysis of the C-terminal pX interactome

The ability of pX to functionally substitute for both the TSG101- and ALIX-interacting late domains of p6^Gag^ in driving EPN release **(Fig. S2A**) suggests it interacts not only with ALIX, but also with other ESCRT components. To assess this, we used label-free quantitative (LFQ) proteomics to analyze proteins precipitated by antibody targeting the Myc tag in EPN from lysates of 293T cells expressing EPN-pX versus EPN-PAD, which lacks the C-terminal 44 amino acid sequence of pX and is not released **(Fig. S5A,B)**. Triplicate samples of the two precipitates were subjected to tryptic digestion and peptide fragments identified by mass spectrometry. A total of 2192 cellular proteins were identified in the two sample sets. Normalized LFQ intensities and sequence coverage were comparable in each sample for peptides derived from the N-terminal pX sequence common to both EPN-pX and EPN-PAD, indicating similar anti-Myc precipitation efficiencies during preparation of the samples **(Fig. S5C,D)**. Eighty-seven cellular proteins were enriched greater than 4-fold in EPN-pX versus EPN-PAD precipitates, suggesting they interact directly or indirectly with the C-terminus of EPN-pX (**Fig. 4A**, **Table S1)**. As expected, these proteins included ALIX. However, the ALIX paralog, HD-PTP (encoded by *PTPN23*), was equally enriched in the complex pulled down with EPN-pX **(Figs. 4A, S5E,F)**. HD-PTP shares structural homology with ALIX, interacts with ESCRT-0 (STAM2) and ESCRT-III (CHMP4B), and is known to contribute to the sorting of ubiquitylated cargo into MVEs [42–44].

**Figure 4.**
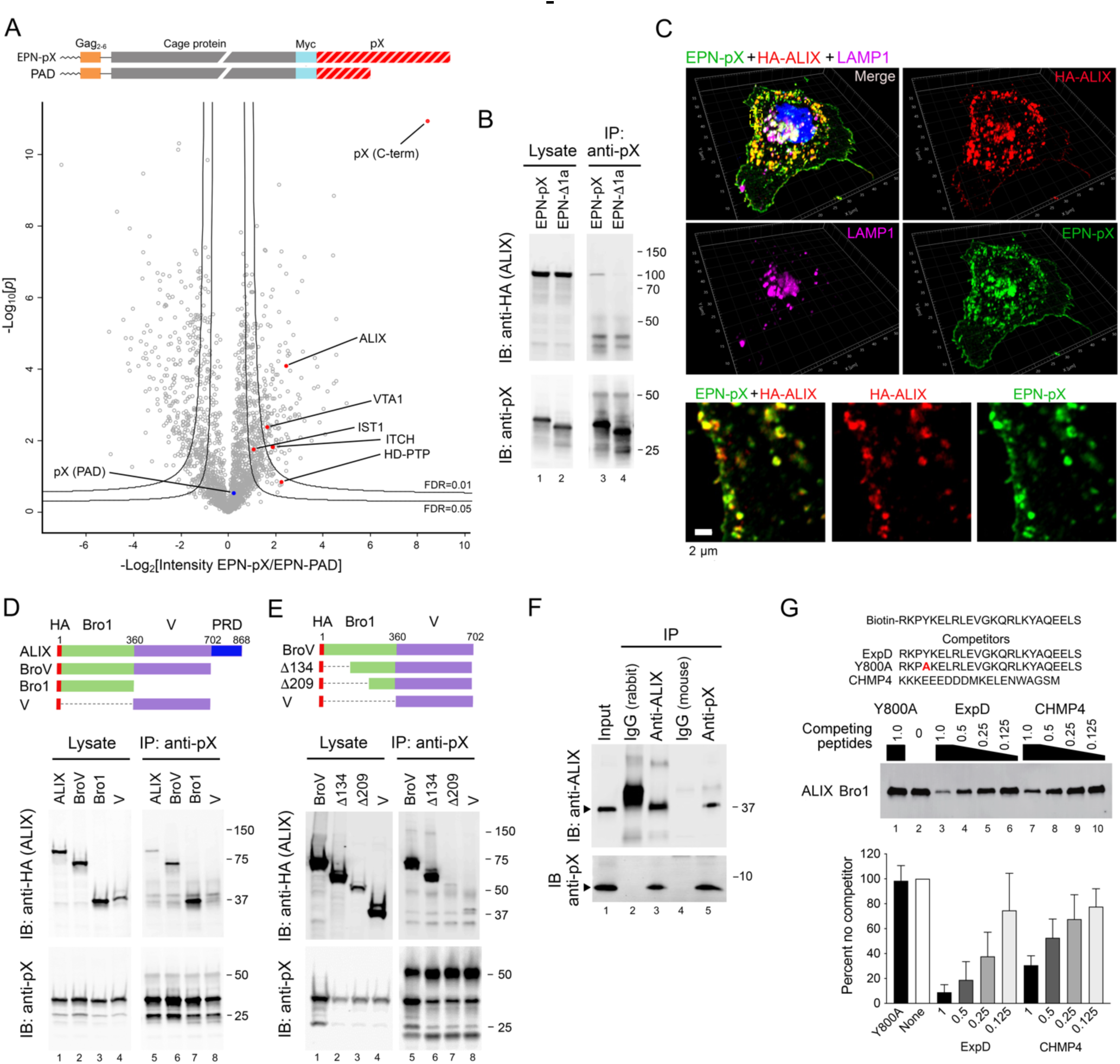
pX interacts directly with the Bro1 domain of ALIX. **(A)** EPN-pX constructs expressed in 293T cells for label-free quantitative (LFQ) proteomic analysis of the pX interactome. Below is a volcano plot showing differential abundance of proteins (LFQ intensities) identified in anti-Myc precipitates of EPN-pX versus EPN-PAD lysates. Differences in normalized LFQ peptide intensity are plotted versus significance (*p* value). Lines represent lower limits of FDR <0.01 and <0.05. Each protein is represented by a single data point with proteins of interest labelled. (**B**) Co-immunoprecipitation of HA-tagged ALIX with EPN-pX. Proteins in lysates of 293T cells transfected with EPN-pX or EPN-Δ1a were immunoprecipitated with anti-pX, followed by blotting with (*top*) anti-HA (ALIX) and (*bottom*) anti-pX. Whole cell lysates are on the left. IP: immunoprecipitation; IB, immunoblot. (**C**) Merged, single-channel Airyscan fluorescent images of a cell transfected with EPN-pX and HA-ALIX expression vectors, showing pX (green), ALIX (HA, red) and LAMP1 (magenta). At the bottom are shown enlarged dual- and single-channel recordings of the region delineated by the yellow rectangle in the merged image at the top. (**D,E)** Co-immunoprecipitation of HA-tagged ALIX or the indicated ALIX domain fragments (*top*) with EPN-pX expressed in 293T cells. **(F)** Co-immunoprecipitation of bacterially-expressed pX and 6XHis-tagged ALIX Bro1 protein (residues 1-392) mixed in buffer containing 0.7% BSA and 0.07% Tween-20. The input mixture (lane 1) was precipitated with the indicated antibodies, followed by immunoblotting as shown. IgG = isotype control. **(G)** Biotin-tagged ExpD peptide pulldown assay. Recombinant ALIX Bro1 protein was incubated with synthetic biotin-tagged ExpD peptide (*top*), affinity isolated on streptavidin beads, and probed with anti-ALIX. Competitor peptides, representing ExpD, the Y800A ExpD mutant, and the C-terminus of CHMP4 [51] (*top*) were added to the mixture at molar ratios decreasing from 1 to 0.125 relative to the biotin-tagged peptide. At the bottom is shown a quantitative analysis of Bro1 pulled down by biotin-tagged ExpD, normalized to pulldown in the absence of competitor peptide. Results shown are means ± s.e.m. from n = 3 independent experiments.

An additional 268 proteins were over 2-fold enriched in EPN-pX precipitates **(Table S1)**. Among these were the ESCRT-III associated proteins IST1 (IST1 homolog), which we identified previously in extracellular eHAV virions and is required for efficient eHAV release [7], and VTA1 (vacuolar protein sorting-associated protein VTA1), which regulates VPS4 ATPase and helps to coordinate disassembly of ESCRT-III complexes [45] **(Figs. 4A, S5E,F)**. Interestingly, the NEDD4 family E3 ubiquitin ligase ITCH (E3 ubiquitin-protein ligase Itchy homolog), which regulates ESCRT recruitment during budding of multiple enveloped viruses [46, 47], was also present in the complex, suggesting it could play a role in EPN-pX release **(Fig. 4A, Fig. S5E,F)**. Consistent with eHAV originating from MVEs, proteins over-represented in the EPN-pX precipitate were highly enriched for components of exosomes and lysosomes **(Fig. S5G)**.

### The ExpD sequence binds the Bro1 domain of ALIX

We confirmed that pX interacts with ALIX by showing that HA-ALIX co-immunoprecipitates with EPN-pX, but not the EPN-Δ1a or EPN-YxxL/A mutants, when co-expressed in 293T cells **(Figs. 4B, S6A)**. Moreover, the degree to which EPN-pX mutants containing Ala or Arg substitutions of conserved ExpD residues co-immunoprecipitated with ALIX correlated well with the impact of these substitutions on EPN-pX release (Spearman r=0.836, **Fig. S6B)**. Super-resolution fluorescence imaging also revealed striking co-localization of ectopically expressed ALIX and EPN-pX in 293T cells, both on internal membranes and at buds on the plasma membrane **(Figs. 4C, S6C)**. By contrast, there was little to no colocalization of ALIX with the EPN-Δ1a mutant **(Fig. S6D)**.

ALIX consists of an N-terminal, boomerang-shaped Bro1 domain, a central ‘V’ domain bent into a V-like conformation, and a C-terminal proline-rich ‘PRD’ domain [15, 48, 49] (**Fig. 4D**). Co-immunoprecipitation experiments with various ALIX deletion mutants indicated that pX interacts with the N-terminal Bro1 domain rather than the V domain, with which the p6^Gag^ YPX_3_L late domain interacts [50] (**Fig. 4D**). These results stand in sharp contrast to those in a previous report that suggested pX interacts with the V domain of ALIX [24]. While deletion of the N-terminal 134 ALIX residues had no effect, further deletion of residues 134-209, which span helices 4-6 near the center of the Bro1 structure [48], ablated the interaction with pX (**Fig. 4E**). To ascertain whether the pX interaction with ALIX is direct or indirect, possibly bridged by other proteins in the complex, we expressed the 8 kDa pX sequence in *E. coli* and purified it to >95% homogeneity. This recombinant pX efficiently formed a complex with a second recombinant protein representing the Bro1 domain of ALIX (residues 1-392, also produced in *E. coli*) that could be precipitated by antibody to either ALIX or pX (**Fig. 4F**). We also synthesized a 24 amino acid peptide representing the ExpD sequence conjugated at its N-terminus to biotin. This biotin-linked ExpD peptide bound the recombinant Bro1, allowing its recovery on streptavidin-coated beads (**Fig. 4G**). Importantly, the biotin-tagged peptide interaction was efficiently blocked by a non-biotin-conjugated ExpD peptide, but not a related peptide with a single Ala substitution at Tyr800 (Y800A) (**Fig. 4G**). Binding was also blocked by a peptide representing the C-terminal 20 residues of the ESCRT-III protein, CHMP4B (**Fig. 4G**), which is known to interact with and contact helices 6-7 of ALIX Bro1 [51]. Taken collectively, these results show the ExpD segment of pX interacts directly with the Bro1 domain of ALIX, overlapping at least partially the CHMP4B binding site. Nonetheless, although pX binds the Bro1 domain of ALIX, overexpression of the entire ALIX protein, including the V and C-terminal PRD domains, was required to enhance EPN-pX release from 293T cells **(Fig. S6E)**.

### pX binds to the Bro1 domain of HD-PTP

HD-PTP is an ALIX paralog with a similar multi-domain structure within its N-terminal 714 residues. It acts non-redundantly with ALIX at endosomes in MVE formation [44, 52]. The N-terminal Bro1 domains of ALIX (residues 5-360) and HD-PTP (residues 10-362) share 32.8% amino acid identity (49.9% similarity). Co-immunoprecipitation experiments with HA-tagged HD-PTP indicated that it forms a complex with EPN-pX when co-expressed in 293T cells, but not with EPN-PAD or EPN-YxxL/A (**Fig. 5A**), confirming the proteomics results shown in **Fig. 4A**. Similar experiments with the EPN-pX mutants containing single amino acid substitutions demonstrated an impact on HD-PTP co-immunoprecipitation similar to the impact on ALIX co-immunoprecipitation, with the exception of the Arg replacement of Ala815 which did not reduce HD-PTP co-immunoprecipitation (**Fig. 5B**, compare with **Fig. S6A)**. As with ALIX, co-immunoprecipitation experiments indicated pX interacts with the Bro1 domain of HD-PTP (**Fig. 5C**). Similarly, the recombinant pX protein was efficiently precipitated with antibody to HD-PTP (and vice versa) when mixed with purified recombinant HD-PTP Bro1 domain expressed in *E. coli* (**Fig. 5D**). We also carried out pulldown experiments using the biotin-tagged ExpD peptide and HD-PTP Bro1 protein expressed *in vitro* in a coupled transcription/translation reaction (**Fig. 5E**). As with ALIX Bro1 (**Fig. 4G**), ExpD peptide pulldown of HD-PTP Bro1 was efficiently blocked by free ExpD and CHMP4 peptides, but not the ExpD Y800A mutant. These results indicate that pX interacts with the Bro1 domain of HD-PTP in a manner similar to its interaction with ALIX. Super-resolution fluorescence imaging was consistent with this, showing that ectopically-expressed HA-tagged HD-PTP colocalized with EPN-pX on the plasma membrane as well on internal membranes of cells (**Figs. 5F,G**).

**Figure 5.**
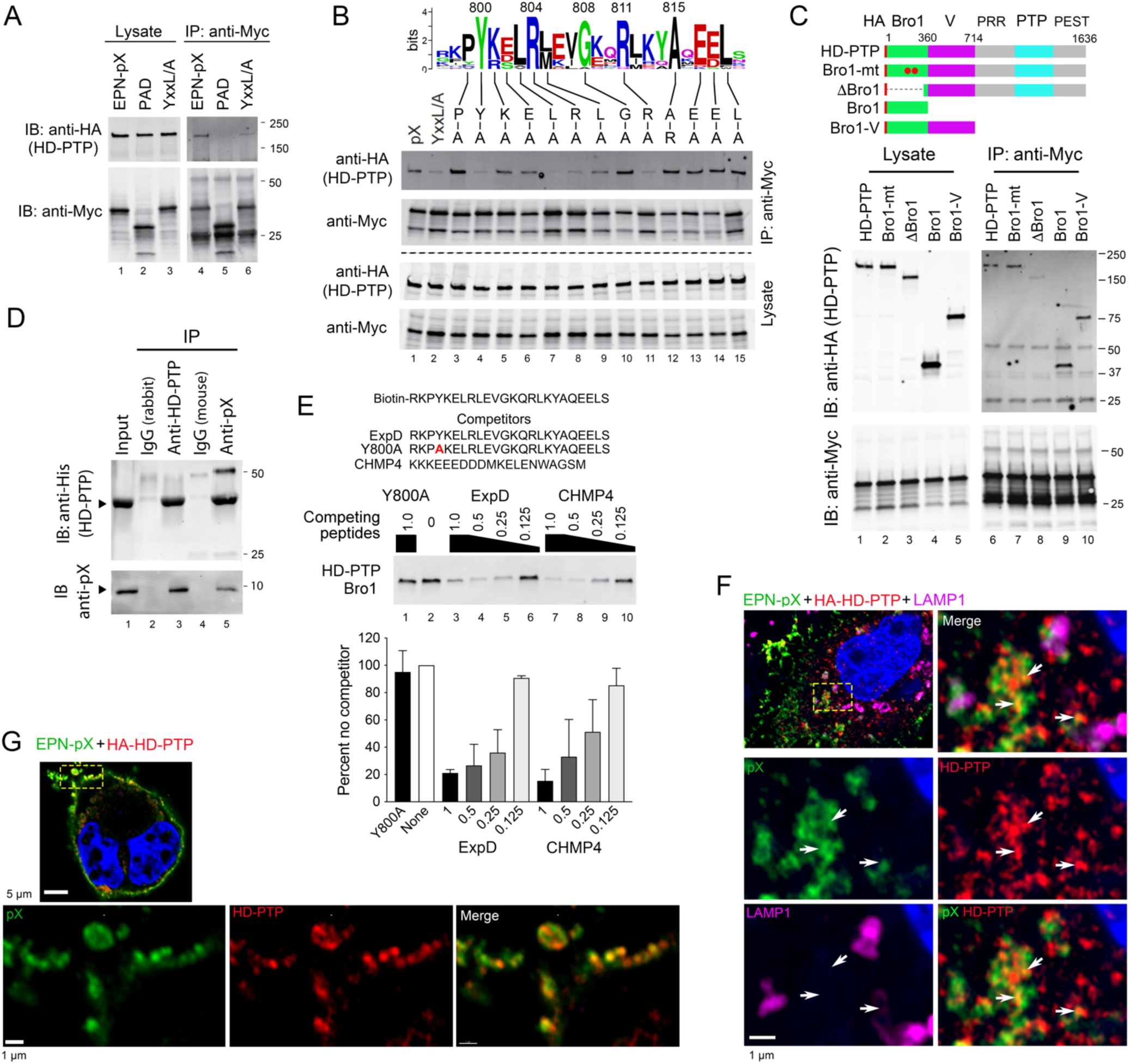
pX interacts directly with the Bro1 domain of the ALIX paralog, HD-PTP. (**A**) Co-immunoprecipitation of HA-tagged HD-PTP with EPN-pX. Proteins in lysates of cells expressing EPN-pX, EPN-PAD (Fig. 2A) or EPN-YxxL/A (Fig. 3A) were precipitated with anti-Myc antibody and immunoblotted with antibody to HD-PTP. **(B)** Co-immunoprecipitation assays of HD-PTP with EPN-pX mutants containing single amino acid substituitions of conserved residues in the ExpD sequence. See legend to Fig. 3A. (**C**) Co-immunoprecipitation assays of EPN-pX with HD-PTP, HD-PTP with L202D and I206D substitutions in the Bro1 domain that eliminate CHMP4 binding (Bro1-mt) [44], and various HD-PTP domain fragments. PRR, proline-rich region; PTP, protein tyrosine phosphatase; PEST, “rich in proline, glutamate, serine and threonine” [52] **(D)** Co-immunoprecipitation of bacterially-expressed pX and recombinant HD-PTP Bro1 protein (residues 1-360) produced in *E. coli*. **(E)** Biotin-tagged ExpD peptide pulldown assay of HD-PTP Bro1 domain (residues 1-360) produced by *in vitro* transcription/translation in rabbit reticulocyte lysate. See legend to Fig. 4G for details. **(F)** Merged low magnification (*top left*) and merged and single-channel enlarged super-resolution Airyscan fluorescence images of the area demarcated by the yellow lines showing pX (green), HD-PTP (HA, red), and LAMP1 (magenta) in a 293T cell expressing EPN-pX and HA-HD-PTP. Arrows indicate sites of pX colocalization with HD-PTP. **(G)** Similar low magnification (top image) and enlarged super-resolution images showing co-localization of pX and HD-PTP along the plasma membrane.

### HD-PTP is required for ESCRT-dependent release of EPN-pX and eHAV

RNAi-mediated depletion of HD-PTP significantly suppressed EPN-pX release from transfected 293T cells (**Fig. 6A**). Release was rescued by overexpression of HD-PTP, confirming the specificity of the knockdown and a requirement for HD-PTP in extracellular release of the nanocage (**Fig. 6B**, lanes 2 versus 3). EPN-pX release was rescued to a lesser extent by overexpression of an HD-PTP mutant (Bro1-mt) with L202D and I206D substitutions in the Bro1 domain that eliminate CHMP4 binding [44], and not at all by a Bro1 deletion mutant (ΔBro1) (**Fig. 6B**, lanes 4 and 5). Interestingly, overexpressing the Bro1 domain only, or Bro1-V (residues 1-714), also rescued EPN-pX release (**Fig. 6B**, lanes 6 and 7). The fact that EPN-pX release is suppressed by depleting either ALIX (**Fig. 3D**) or HD-PTP (**Fig. 6A**) indicates that these Bro1 domain proteins function in a coordinate rather than redundant fashion to mediate release. Nonetheless, overexpression of HD-PTP, or the Bro1 or Bro1-V domains of HD-PTP, substantially rescued EPN-pX release from 293T cells with CRISPR-Cas9 mediated ALIX depletion (ALIX-KO cells) (**Fig. 6C**, lanes 8-14). Unlike ALIX, which is distributed broadly throughout the cell and supports multiple ESCRT-related processes, HD-PTP appears to function selectively at endosomes [44, 52]. The rescue of EPN-pX release from ALIX-KO cells by HD-PTP may have resulted from nonphysiologic localization of the ectopically-expressed protein, as fluorescence microscopy demonstrated its presence at the plasma membrane as well as at internal locations within transfected cells (**Fig. 5G**).

**Figure 6.**
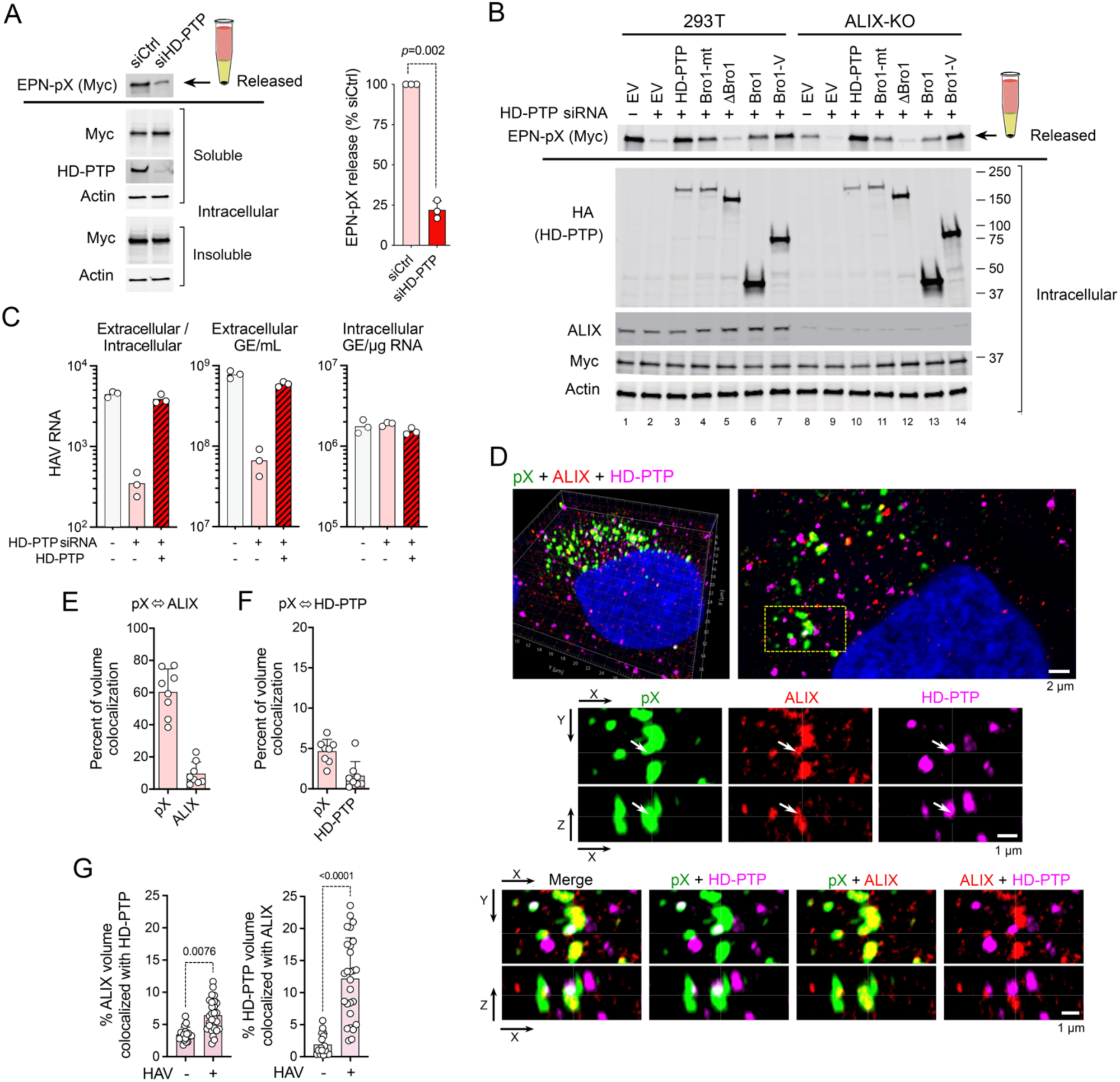
HD-PTP is functionally required for EPN-pX and HAV release. (**A**) EPN-pX release from 293T cells with RNAi-mediated knockdown of HD-PTP (encoded by *PTPN23*) (see legend to Fig. 2D for details of release assay). siCtrl = scrambled siRNA pool. Quantitative analysis on the right with release normalized to release from siCtrl-transfected cells. n=3 independent experiments. *p*-value by t-test with Welch’s correction. **(B)** Rescue of EPN-pX release by overexpression of HD-PTP, HD-PTP with L202D and I206D substitutions (Bro1-mt), or HD-PTP deletion mutants (see Fig. 5C) in HD-PTP-depleted 293T cells (lanes 2-7) and CRISP-Cas9 engineered 293T ALIX knockout (ALIX-KO) cells (lanes 9-14). siRNA: ‘+’ = siHD-PTP, ‘-’ = scrambled siCtrl. EV = empty vector. **(C)** (*left to right*) Ratio of extracellular/intracellular HAV RNA, extracellular HAVl RNA, and intracellular HAV RNA, 48 hrs after cell-free virus infection of Huh-7.5 cells with and without prior RNAi depletion of HD-PTP. Data shown are technical replicates from one of 3 experiments with similar results. GE=genome equivalent. **(D)** Merged and single-channel super-resolution Airyscan fluorescent microscopy images of pX (green, labelled with monoclonal antibody), endogenous ALIX (red), and endogenous HD-PTP (magenta) in an 18f virus-infected Huh-7 cell. A 3D reconstruction is shown on the top left with a sectional view to the right. Below are single- and dual-channel images shown in two projections (X-Y and X-Z). Arrows in the middle panels indicate a prominent site of pX colocalilzation with both ALIX and HD-PTP. **(E)** Quantitative estimates of pX-ALIX volume colocalization in Z-stack images: the first column shows the percentage of pX volume (voxels with pX signal above threshold) containing ALIX signal above threshold; the second column shows the percent ALIX volume containing pX signal. (n=8 cells). See Methods for details. **(F)** Quantitative volume analysis of pX colocalization with HD-PTP similar to that in panel G. **(G)** Voxel colocalization of ALIX with HD-PTP (*left*) and HD-PTP with ALIX (*right*) in HAV-infected and uninfected Huh-7.5 cells. Adjusted p-values determined by ANOVA with Kruksal-Wallis test. n=17-27 cells from multiple experiments.

As with the nanocage protein, HD-PTP depletion significantly reduced the release of virus from infected Huh-7.5 hepatoma cells (**Fig. 6C**). This defect in release was reversed by overexpression of HD-PTP (**Fig. 6C**). The involvement of both ALIX and HD-PTP in virus release was further supported by super-resolution fluorescence microscopy of virus-infected cells, which revealed pX and capsid antigen in close proximity to endogenously expressed ALIX and HD-PTP **(Figs. 6D, S7A,B)**. Quantitative analysis of recorded 3D Z-stack images indicated that 60% ± 5 s.e.m. of the volume occupied by fluorescence signal specific to pX also contained signal above background specific for endogenous ALIX (**Fig. 6E**). Much less colocalization was evident with endogenous HD-PTP, with only 4.7% ± 0.5% of the pX signal sharing the same volume with HD-PTP (**Fig. 6F**). As might be expected, only a minor fraction of either endogenous ALIX (9.3% ± 2.5 s.e.m.) or HD-PTP (1.6 ± 0.6%) colocalized with the viral protein. Colocalization occurred primarily on internal membranes **(Figs. 6D, S7A,B)**, and not on the plasma membrane as observed with ectopically expressed HA-tagged proteins (**Fig. 5G**). Importantly, ALIX fluorescence strongly co-localized with foci of HD-PTP colocalization with pX (*i.e.*, three-way colocalization), and there were highly statistically significant increases in the percentage of ALIX volume colocalizing with HD-PTP, and HD-PTP volume with ALIX, in infected versus uninfected cells (**Fig. 6G**).

## Discussion

The C-terminal pX extension of VP1 found in hepatoviruses is unique among picornaviruses. Although its origins remain obscure, recent crystallographic studies of the downstream 2B protein suggest pX may represent the vestigial N-terminal remnant of a 2A protease present in an enterovirus-like ancestor [53]. In this report, we show that pX contains crucial determinants of quasi-envelopment, acting as an adaptor that recruits ESCRT to the viral capsid by binding two distinct Bro1-domain proteins, ALIX and HD-PTP. This adaptor function is central to the biology of the virus and conserved among pX sequences of hepatoviruses infecting mammalian species separated by tens of millions of years evolution (**Fig. 3B**).

ALIX and HD-PTP are structurally and functionally related [25, 42]. Both bind CHMP4B within their N-terminal Bro1 domains and serve as scaffolds for assembly of ESCRT-III complexes capable of sculpting membranes. However, ALIX and HD-PTP differ in how they are regulated and where they function within the cell. ALIX is broadly expressed and involved in multiple membrane abscission events including cytokinesis, nuclear envelope reformation, endolysosomal repair, and MVE formation, as well as budding of canonical enveloped viruses [54, 55]. By contrast, HD-PTP functions more selectively at endosomes and in MVE formation [25, 42, 56]. It directs the sorting of activated epidermal growth factor receptor (EGFR) into intralumenal vesicles during MVE formation, a process in which ALIX has little or no role [42, 44]. Although over-expression of the isolated HD-PTP Bro1 domain boosts release of virus-like particles formed by a minimal HIV-1 Gag protein [57], HD-PTP has not been implicated previously in the release of any type of virus from cells.

The self-assembling nanocage protein [29] provided a useful model that recapitulates the role of pX in viral egress, allowing for pX expression levels much higher than possible with infectious virus and facilitating functional evaluation of pX mutants. Experiments with EPN-pX constructs, coupled with a phylogenetic analysis of pX sequences identified a conserved peptide motif, YxxLR[L/M]xxGxxRxxxA (**Figs. 3**, **Fig. S3)**, that is crucial for binding to the Bro1-domain proteins and for nanocage release. The HIV-1 p6^Gag^ protein contains two distinct late domains, one of which (PTAP) binds TSG101, and the other (YPX_3_L), binds ALIX. This provides links to both ESCRT-I and ESCRT-III. Both late domains are required for release of nanocages fused to p6^Gag^ [17] **(Fig. S2A)**. By contrast, the 24 amino acid ExpD sequence functioned well as a single determinant in driving EPN-pX release (**Fig. 3C-E**). This difference may arise from the capacity of pX to recruit HD-PTP as well as ALIX, as well as the manner in which pX interacts with these Bro1 domain proteins.

Jiang and colleagues [24] reported that fusing pX to eGFP resulted in ALIX-dependent sorting of eGFP into MVEs and the release of eGFP in exosomes. They suggested pX interacts with the V domain of ALIX in a manner similar to the YPX_3_L motif of lentiviruses [58]. In contrast, our data indicate pX interacts directly with the Bro1 domain of ALIX, with residues 134-209 being crucial for the interaction (**Fig. 4D-F**). pX also interacts with the related N-terminal Bro1 domain of HD-PTP (**Fig. 5C-E**). Unlike ALIX, the Bro1 domain of HD-PTP binds STAM2, a key component of ESCRT-0, in addition to CHMP4, the core component of ESCRT-III that is bound by both ALIX and HD-PTP [43]. By binding HD-PTP, pX may recruit ESCRT-0, functioning like the PTAP late domain of HIV-1 p6^Gag^ that interacts with ESCRT-I by binding TSG101 [14, 17]. A second difference exists in the basal activation states of ALIX and HD-PTP. Unlike ALIX, which exists basally in a closed, inactive conformation with its V domain occluding the Bro1 CHMP4-binding site, HD-PTP assumes an open, linear conformation in which its Bro1 CHMP4- and STAM2-binding sites are freely available for interaction [52]. HD-PTP also differs from ALIX in its extended carboxy-terminal protein tyrosine phosphatase (PTP) and PEST domains [52, 59]. However, neither of these domains were critical for restoration of EPN-pX release from HD-PTP-depleted cells (**Fig. 6B**).

The structural details of the pX-Bro1 interaction remain to be elucidated. Cryo-electron microscopy of purified EPN-pX nanocages showed no clearly discernable density for pX (**Fig. S1C**). However, we did observe significant decoration adjacent to the C-terminus of the nanocage protein, of the correct volume for pX, in a structure reconstructed from data collected on EPN-pX nanocages released by spontaneous lysis of EPN-filled vesicles during blotting and vitrification (**Fig. S1D,E**). It seems likely that surface tension during blotting drove vesicle lysis and rapid vitrification then allowed intact EPN-pX nanocages to be imaged and particles bearing pX to be reconstructed to low resolution. The pX domain appears suitably positioned to interact with the Bro1 proteins and other ESCRT components identified in the proteomics study. A peptide representing the C-terminus of CHMP4 competed efficiently with binding of the ExpD peptide to both ALIX and HD-PTP (**Fig. 4G**, **5E**). This suggests the likelihood that pX interacts with a hydrophobic patch on the surface of Bro1 formed by residues between Phe-199 and Leu-216 in ALIX that is recognized by both CHMP4 and the Y3 protein of Sendai virus [51, 60].

The efficient release of quasi-enveloped virus and EPN-pX requires both ALIX and HD-PTP (**Fig. 3D**, **6A-C)** [1]. This indicates that ALIX and HD-PTP do not function redundantly in parallel pathways, but rather suggests a model in which these proteins act in a coordinated fashion during the egress of eHAV, perhaps facilitated by the 60 copies of pX displayed on the capsid surface (**Fig. 7**). Further experiments are needed to fully understand this, but it seems likely that the requirement for HD-PTP reflects the egress of eHAV through the endosomal MVE pathway, rather than release from the plasma membrane of the cell. This would be consistent with proteomics studies showing eHAV vesicles are highly enriched in MVE-associated proteins [7]. The constituitvely open activation state of HD-PTP might also account for its requirement in eHAV release. In addition to the Bro1 proteins, we identified the E3 ubiqutin ligase ITCH among proteins associating with EPN-pX (**Fig. 4A**). Although a role in eHAV release remains to be studied, NEDD4-family E3 ligases like ITCH are involved in the budding of many conventional enveloped viruses, including HIV-1 [47, 61, 62]. Ubiquitylation is a common signal for cargo to be trafficked into MVE, and the V domains of both ALIX and HD-PTP have specific ubiquitin-binding activities [42, 63, 64].

**Figure 7.**
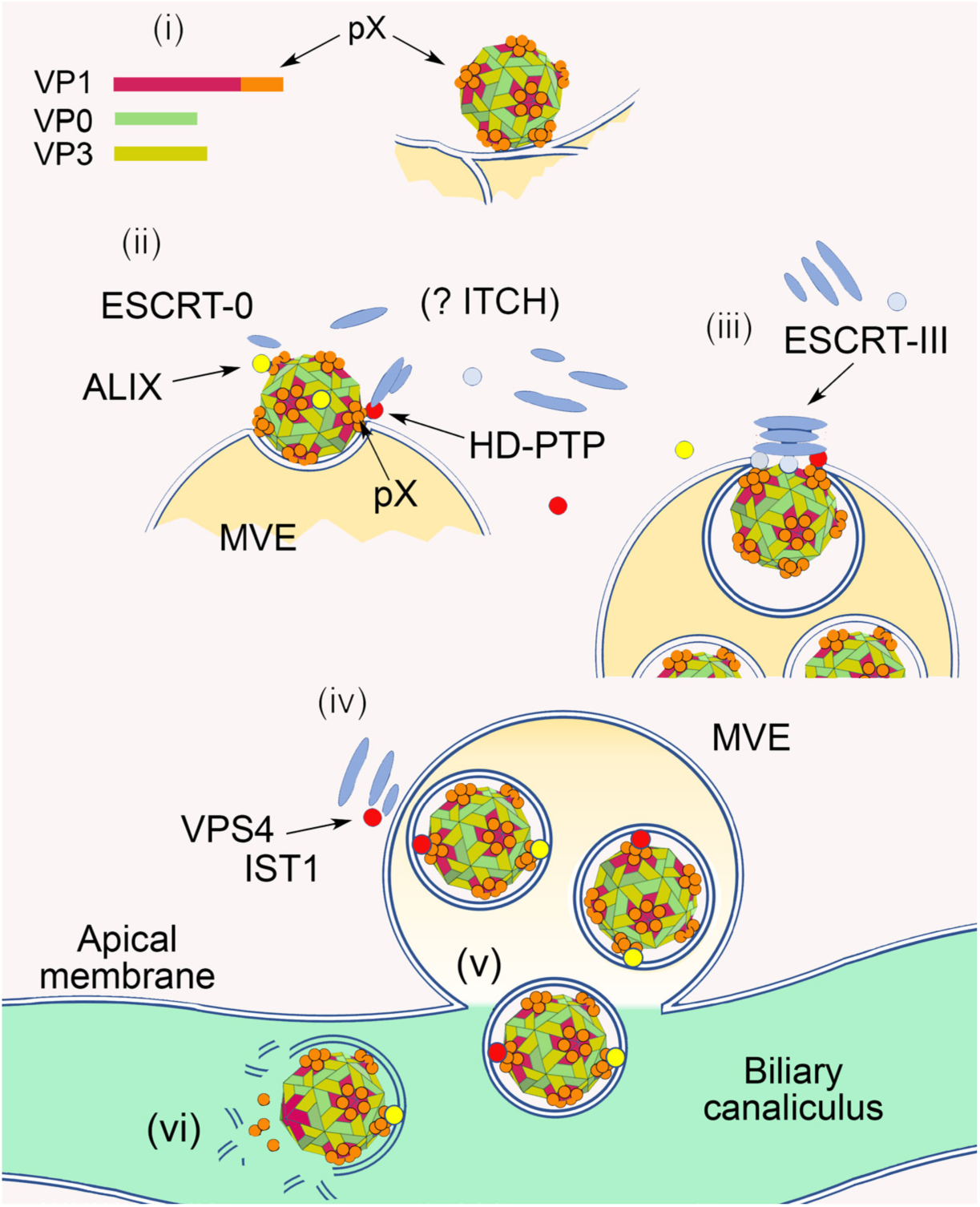
The Bro1-domain proteins HD-PTP and ALIX function non-redundantly in nonlytic hepatovirus release from infected hepatocytes. (i) Assembly of membrane-associated capsids decorated on their surface with 60 copies of pX. (ii) pX recruits both ALIX and HD-PTP to capsids. (iii) HD-PTP and ALIX act coordinately to recruit STAM2 (ESCRT-0) and CHMP4, scaffolding assembly of an ESCRT-III complex that facilitates inward budding of the capsid on an endosome and mediates abscission of the membrane leading to formation of a multivesicular endosome (MVE). (iv) Upon membrane abscission, VPS4 mediates disassembly of ESCRT-III, regulated in part by IST1, VTA1 and HD-PTP. (v) MVEs traffic to the apical plasma membrane of the hepatocyte where membrane fusion releases quasi-enveloped eHAV into the biliary canaliculus. (vi) Bile salts degrade the eHAV membrane resulting in naked nHAV particles being released from the biliary tract into the small intestine [10].

The nonlytic release of quasi-enveloped eHAV distinguishes hepatoviruses from other genera in the family *Picornaviridae* [1]. Other picornaviruses, including both enteroviruses and cardioviruses, are released from infected cell cultures in EVs of various sizes [2–4], but *in vivo* data showing a primary role for such release in pathogenesis exist only for hepatoviruses [1]. The ESCRT adaptor function of pX, coupled with YPX_3_L late domains identified previously in the VP2 capsid protein [1, 16], also distinguish hepatoviruses from other picornaviruses for which there is as yet no evidence for late domains capable of recruiting ESCRT. pX represents a specific evolutionary adaptation that allows HAV, a prototypically nonenveloped picornavirus, to be released from cells in the absence of cell death, sequestering it from the host immune system and promoting its spread *in vivo* to uninfected cells.

## Materials and Methods

### Cells and virus

H1 Hela and human hepatocyte-derived Huh-7.5 [65] cells were mycoplasma-free and cultured as described previously [66]. HEK293T (293T) cells were procured from the ATCC (CRL-3216). Viruses used in these studies were derived from modified infectious molecular clones of low or high cell culture-passage HM175 strain HAV, pT7-p16.2 (GenBank KP879217.1) and pT7-18f.2 (KP879216.1), respectively, containing 40-nt long 3’ poly-A tails [67, 68]. The 18f-NLuc reporter virus expressing nanoluciferase has been described previously [69].

### Plasmids and oligonucleotides

Plasmids encoding *de novo* designed protein nanocages fused to wildtype or mutant HIV p6^Gag^ sequence (EPN01N-based vectors) and VPS4A(E228Q) [29, 30], ALIX [50], and expression vectors for *Gaussia princeps* luciferase fragments [27] have been described previously. pHA-IST1 (#131619), pRK5-HA-Ubiquitin-WT (#17608) and pRK5-HA-Ubiquitin-K0 (#17603) were from Addgene. Vectors expressing the nanocage fused to wild-type or mutant pX sequences, ALIX mutants, and GLuc2-VP2 mutants were constructed using PCR-based strategies. DNA encoding a nanocage fragment fused to pX sequences of *Eidolon helvum* M32 virus (GenBank KT452714.1) and *Miniopterus manavi* SMG18520 virus (KT452742.1) was synthesized by Genewiz (South Plainfield, NJ) and ligated into pEPN-pX at *EcoR*1 and *Afl*2 sites to produce pEPN-M32pX and pEPN-SMGpX, respectively. GLuc2-VP1pX, -VP1 and -pX were constructed using Gateway cloning. pT7-16.2 mutants with Ala and Arg substitutions in pX were constructed by splicing synthetic cDNA from Genewiz into *Sac*1 and *PflM*1 sites. All final constructs were verified by DNA sequencing. Oligonucleotides used in this study are listed in Table S2.

### Antibodies

A murine monoclonal antibody to pX (clone 59.7.1) was raised against bacterially-expressed 6xHis-tagged HM175 pX protein (RRID:AB_2868525, 1:1000). Other antibodies used in this study are listed in Table S3. Species-specific Alexa Fluor conjugated secondary antibodies were purchased from Thermo Fisher Scientific.

### Chemical reagents

IMP-1088 was purchased from Cayman Chemical (Cat# 25366-1). Iodixanol (Opti-Prep Density Gradient Medium) was purchased from Millipore-Sigma (Cat# D1556). SMARTPool ON-TARGETplus and non-targeting control #2 siRNAs were purchased from Dharmacon.

### siRNA and CRISPR/Cas9 protein depletion

RNAi depletion of cellular proteins and lentivirus-transduced CRISPR/Cas9 knockout of protein expression were carried out as described previously [69]

### *In vitro* transcription and RNA and DNA transfections

Genome-length RNA (5 µg) was transcribed *in vitro* from plasmid DNA and electroporated into 5×10^6^ Huh-7.5 cells as described previously to generate virus [70]. DNA transfections were carried out with FuGENE HD (Promega) according to the manufacturer’s recommended protocol. Briefly, 293T cells (5×10^5^ per well in 6-well plates) were transfected with 500 ng of EPN vectors, or co-transfected with an equal amount of empty or VPS4A(E228Q) expression vector. For co-immunoprecipitation experiments, 2.5×10^6^ 293T cells, seeded previously in 10-cm dishes, were co-transfected with 500 ng EPN vector and 5 μg empty or HA-tagged ALIX, or HD-PTP expression vectors. For proteomic analysis, similarly seeded 293T cells were transfected with 2.5 μg EPN-pX or EPN-PAD vector. The Lipofectamine RNAiMAX transfection reagent (Thermo Fisher Scientific) was used for transfection of 50 nM SMARTPool ON-TARGETplus or #2 control siRNAs (Dharmacon), according to the manufacturer’s recommended procedure.

### Isopycnic gradient centrifugation of virus and measurement of viral RNA

Isopycnic gradient analysis of intracellular virus and virus released into cell culture supernatant fluids was carried out as previously described [1, 16]. Fractions were collected manually from the top of the gradient. RNA was extracted from fractions using the RNeasy kit (Qiagen), followed by two-step RT-qPCR quantitation of HAV RNA.

### Protein-fragment complementation assay

The *Gaussia princeps* luciferase protein-fragment complementation assay was carried out as described previously [27, 28]. Each interaction pair was assessed in at least two independent experiments, each with 3 technical replicates. Positive bait-prey protein interactions were identified by luciferase activities exceeding those obtained with a large panel of control prey proteins [27].

### Nanocage release assay

Release was assayed as previously described [30]. Supernatant fluids were collected from 293T cells 24 hrs after transfection with nanocage vectors, clarified by centrifugation at 8,000 rpm for 10 min, and then centrifuged through a 200 µL 20% sucrose cushion in an Eppendorf 5424R centrifuge at full speed for 90 min. Samples were carefully removed from the bottom of the cushion and suspended in 50 µL Laemmli sample buffer. Cells were lysed in 200 µL lysis buffer (50 mM Tris-pH 7.4, 150 mM NaCl, 1% Triton X-100, and 1x complete protease inhibitor cocktail (Millipore-Sigma) on ice for 5 min. Soluble and insoluble proteins were separated by centrifugation at 12000 rpm for 5 min in an Eppendorf 5424R centrifuge. Released nanocages, and soluble and insoluble intracellular protein fractions, were subjected to SDS-PAGE followed by immunoblotting with anti-Myc antibody. The efficiency of release was calculated as the percentage of EPN protein pelleted from culture supernatant against total EPN protein expressed (normalized intracellar EPN protein plus supernatant fluid EPN protein) based on quantitation of immunoblots using the Odyssey Imaging System (LI-COR Biosciences) .

### Immunoblots and quantification of nanocage release

Immunoblot analysis was carried out using standard methods. An Odyssey Imaging System (LI-COR Biosciences) was used for visualization and signal (infrared fluorescence) intensity analysis.

### Co-immunoprecipitation analysis

Pierce Protein A/G Agarose and magnetic Dynabeads^TM^ Protein G beads (both from Thermo Scientific) were used for co-immunoprecipitation and protein pull-down proteomic analysis respectively. Cell lysates were prepared from 10-cm dish cultures with lysis buffer (20 mM Tris-HCl, pH 7.5, 50 mM KCl, 250 mM NaCl, 10% glycerol, 5 mM EDTA, 1× complete protease inhibitor and 1× PhoSTOP, 0.5% NP-40). The clarified lysates were then used for immunoprecipitation according to the manufacturer’s protocols.

### Nanoparticle tracking analysis

The size and concentration of EVs released by 293T cells were monitored by laser scattering video microscopy using a ZetaView device (Particle Metrix) with a particle cut-off size of 1000 nm [71]. Cells were propagated in media supplemented with exosome-free fetal calf serum, and supernatant fluids clarified by low-speed centrifugation prior to analysis.

### Purification of engineered protein nanocages (EPNs)

Ten 175 cm^2^ flasks of HEK293T cells were transfected with pEPN-pX using polyethylenimine [72]. At 48 hrs, clarified culture medium (10,000 g, 10 minutes) was centrifuged at 150,000 g (r_max_) for 1 hr at 12 °C and the pellet was resuspended in 12 mL PBS. The sample was centrifuged again at 150,000 g (r_max_) for 1 hr at 12 °C and resuspended in 200 µL PBS. It was then loaded on a 5-35 % iodixanol density gradient in PBS and centrifuged at 300,000 g (r_max_) for 4 hrs at 12 °C. The diffuse band of EPNs in the central region of the gradient was collected and diluted in PBS before being centrifuged at 150,000 g (r_max_) for 2 hrs at 12 °C. The pellet was resuspended in 20 µL PBS and stored at 4 °C.

### Cryo-EM sample preparation, data collection and processing

The EPN suspension was applied to glow-discharged copper support R2/2 holey carbon TEM grids (Quantifoil) and vitrified in liquid ethane/propane using a Cryoplunge 3 (Gatan) with a blot time of 4-5 s at >90 % humidity. Grids were screened using a Glacios TEM operating at 200 keV and a data set of 1803 movies was acquired through EPU (Thermo Fisher FEI) at a nominal magnification of 92 kX and a nominal defocus range of -0.7 to -3 µm using a Falcon-III detector (ThermoFisher FEI) in integrating mode. Each acquired image had a total dose of 2.3 e^−^/Å^2^ distributed over 20 frames. Movies were aligned and summed using MotionCor2 [73] before CTF correction using CTFFIND4 [74] on non-dose-weighted image sums. The resulting corrected micrographs were then processed using Relion3.0 [75]. Promising particles were manually selected through the Relion3.0 GUI and extracted particles were filtered by classification with Relion3.0 (1830 particles). Three-dimensional reconstruction was performed using a symmetry-free *ab initio* model derived using Relion’s Initial Model module as a reference with icosahedral symmetry. The final three-dimensional map was masked and B-factor applied via the post-processing module in Relion. A calibrated pixel size was derived from fitting of the previously solved nanocage structure (PDB: 5KPN) (*27*) and determined as 1.51 Å/pix.

### Label-free quantitative proteomics

Lysates were prepared from three independent 10-cm dish cultures of 293T cells transfected with pEPN-pX and pEPN-PAD with lysis buffer (20 mM Tris-HCl, pH 7.5, 50 mM KCl, 250 mM NaCl, 10% glycerol, 5 mM EDTA, 1× complete protease inhibitor and 1× PhoSTOP, 0.5% NP-40) followed by immunoprecipitation with anti-Myc antibody coupled to magnetic beads (Thermo Scientific). The protein samples were separated by SDS-PAGE, extracted from segments cut from the gel, destained, reduced and alkylated and then subjected to tryptic digestion. The peptides were extracted and desalted on home-made C18 stage-tips, then dissolved in 0.1% formic acid and analyzed on a Q-Exactive HF-X coupled with an Easy nanoLC 1200 (Thermo Fisher Scientific, San Jose, CA). Peptides were loaded onto a nanoEase MZ HSS T3 Column (100Å, 1.8 µm, 75 µm x 150 mm, Waters). Analytical separation of all peptides was achieved with 45-min gradient. A linear gradient of 5 to 30% buffer B over 29 min and 30% to 45% buffer B over 6 min was executed at a 300 nl/min flow rate, followed by a ramp to 100% buffer B in 1 min and a 9-min wash with 100% buffer B, where buffer A was aqueous 0.1% formic acid, and buffer B was 80% acetonitrile and 0.1% formic acid. LC-MS experiments were also carried out in a data-dependent mode with full MS (externally calibrated to a mass accuracy of <5 ppm and a resolution of 60,000 at *m*/*z* 200) followed by high energy collision-activated dissociation-MS/MS of the top 15 most intense ions with a resolution of 15,000 at *m/z* 200. High energy collision-activated dissociation-MS/MS was used to dissociate peptides at a normalized collision energy of 27 eV in the presence of nitrogen bath gas atoms. Dynamic exclusion was 20.0 seconds. Each of the three biological replicates was subjected to two replicate technical LC-MS analyses.

### Raw proteomics data processing and analysis

Mass spectra were processed, and peptide identification carried out using MaxQuant software version 1.6.10.43 (Max Planck Institute, Germany). Protein database searches were against the UniProt human protein sequence database (UP000005640), EPN proteein and HAV pX sequences. A false discovery rate (FDR) for both peptide-spectrum match (PSM) and protein assignment was set at 1%. Search parameters included up to two missed cleavages at Lys/Arg on the sequence, oxidation of methionine, and protein N-terminal acetylation as a dynamic modification. Carbamidomethylation of cysteine residues was considered as a static modification. Peptide identifications were reported by filtering of reverse and contaminant entries and assigning to their leading razor protein. The label-free quantitation (LFQ) was carried out using MaxQuant. Data processing and statistical analysis were performed on Perseus (Version 1.6.0.7). Protein quantitation of technical replicates, and two-sample t-test statistics with a threshold p-value of 0.01, were used to report statistically significant protein abundance fold-changes. The mass spectrometry proteomics data have been deposited to the ProteomeXchange Consortium via the PRIDE partner repository with the dataset identifier PXD022107.

### Super-resolution Airyscan fluorescence microscopy

DNA-transfected 293T or 18f-NLuc virus-infected Huh-7.5 cells were visualized in 35mm glass bottom dishes with 14mm bottom wells (Cellvis, Cat# D35-14-1.5-N). Cells were fixed with 4% paraformaldehyde for 12 mins, washed twice with PBS and incubated with a blocking/permeabilization solution containing 5% goat serum and 0.1% saponin in PBS for 1 hr at room temperature. Cells were stained for 1 hr at room temperature with primary antibody diluted in PBS with 1% BSA and 0.1% saponin, then washed with PBS containing 0.05% saponin followed by staining with appropriate secondary antibodies for 1 hr. Nuclei were counterstained with Hoechst 33342 (Invitrogen) for 10 mins at room temperature. Slides were then washed three times with PBS containing 0.05% saponin, and where appropriate incubated for 3 min with a 1:2000 dilution of CellMask Deep Red (Thermo Fisher) in PBS. Imaging data were recorded in super-resolution mode on a laser-scanning confocal Zeiss 880 microscope (Carl Zeiss AG, Oberkochen, Germany) equipped with an Airyscan detector and controlled by Zen Blue 3.0 software. Airyscan super-resolution recording enhances resolution in all directions by a factor of 1.7. The microscope was mounted on a Zeiss Inverted Axioobserver Z1 base equipped with a Definite Focus unit to ensure focus plane stability. The Plan-Apochromat 63X/1.40 Oil DIC M27 objective (Zeiss) was used for all imaging. For each image plane, fluorescence signals were recorded sequentially at different wavelengths in super-resolution mode using appropriate laser excitation and filter sets: for DAPI/ Hoechst, excitation 405nm, emission filter BP 420-480 + BP 495-550; for Alexa488, excitation 488nm, emission filter BP 420-480 + BP 495-550; for Alexa568/594, excitation 561nm, emission filter BP 420-480 + BP 495-620; and, for Alexa647, excitation 633nm, emission filter BP 570-620 + LP 645. Fields of view were selected randomly maintaining identical imaging conditions for different cells in each experiment. The number of X-Y pixels on the frame was set to “optimal” in the Zeiss software controller, with XY pixel size 0.043 µm. The distance between images in the Z direction during Z-stack recording was 0.160 µm. Immediately after recording, the raw data was processed by Zen Blue software (Zeiss) to produce a super-resolution image with 16-bit depth.

### Post-imaging analysis

Imaris software (version 9.6, Bitplane, Zurich) was used for 3D visualization of recorded Airyscan images and to analyze colocalization of fluorescent signals. Following background subtraction, the Imaris colocalization module was used to identify the percentage of 3D image volume within which fluorescence signals were colocalized. Intensity thresholds of fluorescence signals were selected for each fluorescence channel separately using a semi-automatic procedure with the condition that the threshold should not be less than 10% of maximum channel intensity. Separate channels were created for colocalization results obtained from the analysis of each pair of fluorescent signals (*e.g.*, pX or HAV with HD-PTP; and pX or HAV with ALIX). The percentage volume colocalization was then determined for these newly created channels and the fluorescence signal from a third channel, allowing identification of colocalization of all three signals above threshold. The positions and shapes of structures labeled with specific antibodies were visualized by rendering the surfaces of 3D fluorescent signals based on their intensities.

### Statistical analysis

Unless indicated otherwise, significance was assessed by unpaired t test or ANOVA. All statistical calculations were carried out using with Prism 8.4.3 software (GraphPad). Significance values are shown as ****p<0.0001, ***p<0.001, **p<0.01, *p<0.05.

## Acknowledgements

The authors gratefully acknowledge Wesley Sundquist and Jurg Voteller of the University of Utah for their gift of the EPN-p6^Gag^ (EPN01N) and VPS4A(E228Q) plasmids, and for helpful advice and discussions in early stages of the project; Phillip Woodman and Lydia Wunderley of the University of Manchester for HD-PTP expression vectors; and, Dirk Dittmer, UNC Department of Microbiology and Immunology, for assistance with nanoparticle tracking analysis.

## Author contributions

Conceptualization: HF, TShir, SML, SE, EF, DIS

Methodology: HF, TShir, KLM, LX, LW, WF, JAD, RB-B, SK, AH-W

Investigation: HF, TShir, HMED, MB, LS, RM, TShio, LX, LC, RB-B, SK

Visualization: HMED, DIS, MK

Supervision: SML, EF, DIS, SE, XC

Writing - original draft: HF, TShir, SML

Writing - review and editing: SML and all authors

## Data and Materials Availability

Mass spectrometry proteomics data have been deposited to the ProteomeXchange Consortium via the PRIDE partner repository with the dataset identifier PXD022107. All other data are available in the manuscript or the accompanying Supplementary Materials.

## SUPPLEMENTARY MATERIALS

**Figure S1.**
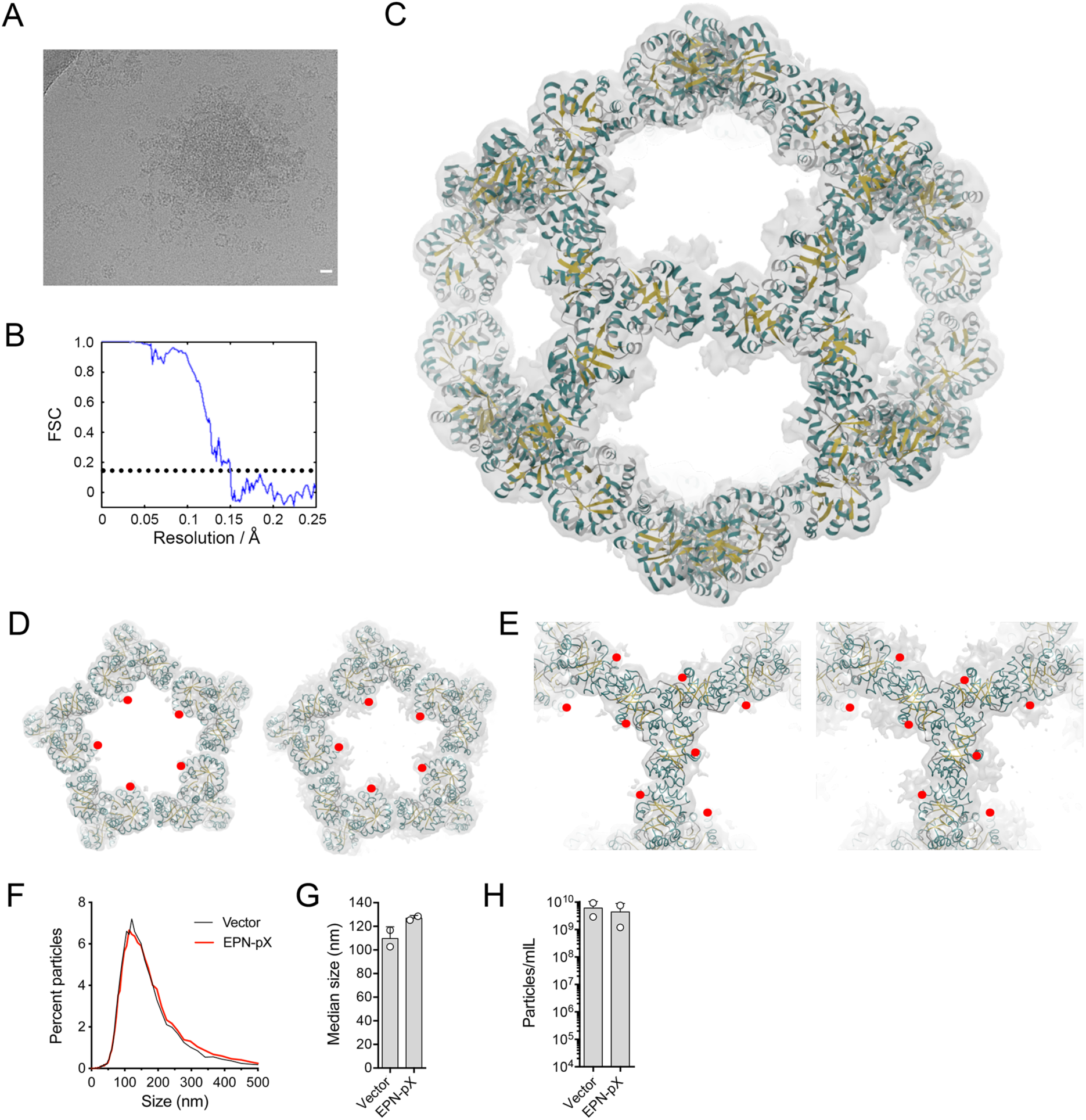
Low-resolution cryo-EM structure of EPN-pX. **(A)** Example electron micrograph of extracellular fluids showing a burst vesicle. Scale bar = 200 Å. **(B)** Fourier shell correlation (FSC) plot showing attained resolution of 6.7 Å. **(C)** Overview of three-dimensional map with a rigid-body fitted to a previously published model of the EPN nanocage (PDB: 5KP9) in cartoon representation. Secondary structural elements are colored with helices in cyan and β-sheets in yellow. **(D)** Exterior views of the five-fold cage face at (*left*) high and (*right*) low contour levels showing additional density that may be consistent with pX. The C-terminus of the EPN sequence is indicated by the red dot. **(E)** Interior views of the three-fold nanocage axis, displayed as in panel D. **(F)** Laser-scattering video microscopy (NTA) estimates of the size distribution of extracellular particles in supernatant fluids of 293T cells 24 hr after transfection with EPN-pX (n=22,114 particles) or empty vector (n=18,038). **(G)** Median particle size measured by NTA in 2 independent transfection experiments. **(H)** NTA estimates of EV concentration in extracellular fluids.

**Figure S2.**
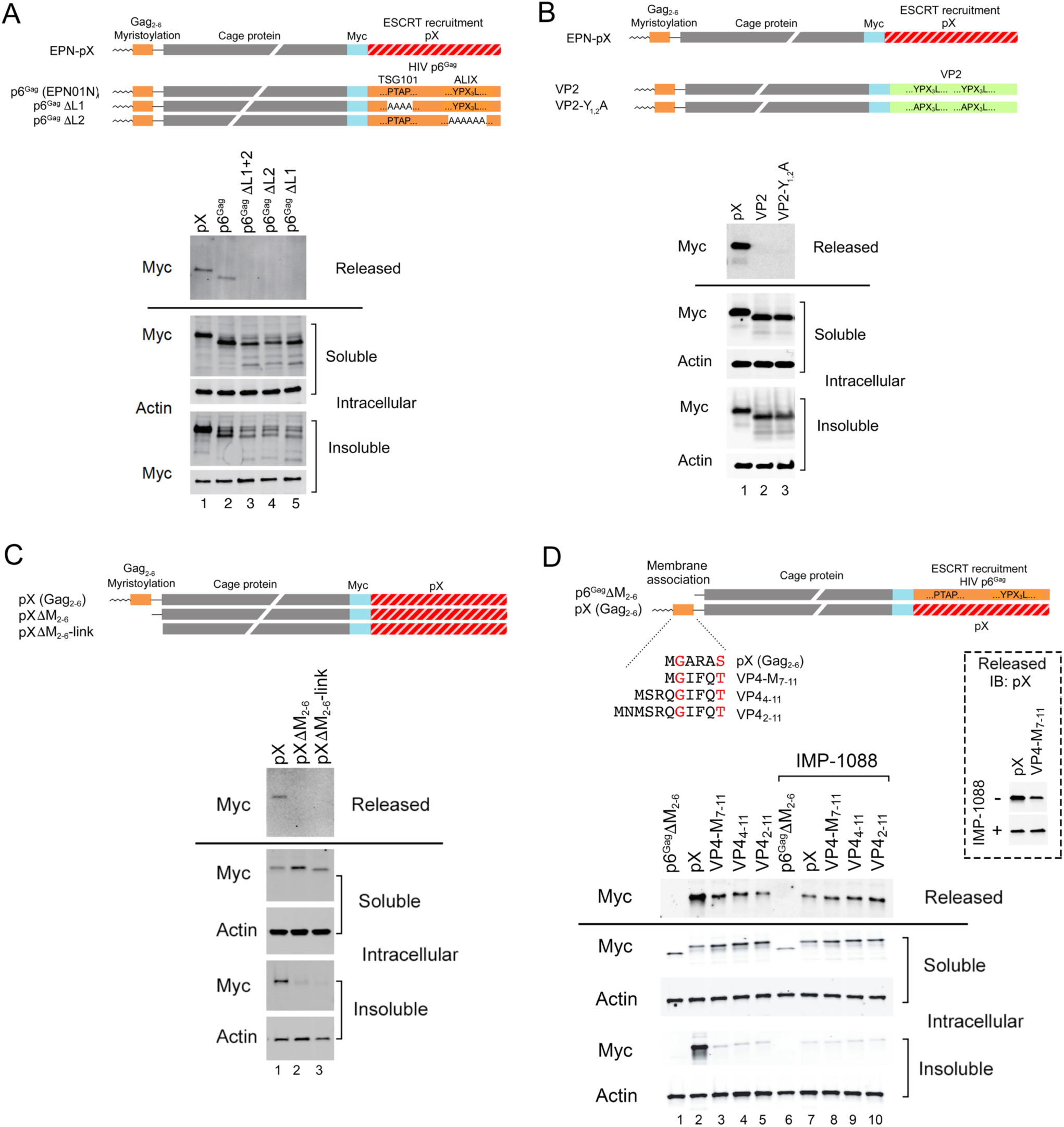
Cellular release of EPN-pX, EPN-p6^Gag^ (EPN01N, reference *29*) and EPN-VP2 constructs. (**A**) (*top*) EPN-pX and EPN-p6^Gag^ constructs with single and dual deletions of the p6^Gag^ late domains that bind TSG101 (PTAP) or ALIX (YPX_3_L) (p6^Gag^Δ1 and p6^Gag^ΔL2, respectively). (*bottom*) Immunoblots showing extracellular release of EPN-pX compared with EPN-p6^Gag^ with and without single or dual deletions of the p6^Gag^ late domains. Deletion of either late domain ablates nanocage release, as shown previously (*29*). (**B**) Absence of detectable extracellular release of EPN-VP2 which contains VP2 residues 130-195 fused to the EPN C-terminus, with and without Ala substitutions of the leading Tyr residue in both putative ALIX-interacting VP2 late domains (VP2-Y_1,2_A) (*16*). EPN-pX was included as a positive control. (**C**) (*top*) EPN-pX nanocage constructs with and without the N-terminal HIV Gag myristoylation signal sequence. The ΔM_2-6_ and ΔM_2-6_-link constructs lack the p6^Gag^ myristoylation signal (M_2-6_) that directs membrane association, while ΔM_2-6_-link also lacks a downstream linker sequence. (*bottom*) Myc immunoblots showing nanocage proteins released into extracellular fluids from transfected 293T cells (recovered after pelleting through a 20% sucrose cushion) and intracellular soluble and insoluble nanocage proteins expressed by the cells. Actin is shown as a loading control. (**D**) (*top*) EPN-pX constructs in which the Gag myristoylation signal (Gag residues 2-6) is replaced with peptide sequences from HAV VP4 (residues numbered according to the HAV ORF). (*bottom*) Myc immunoblots as in panel A. Lanes 6-10 show proteins released/expressed in cells treated with the N-myristoylation inhibitor IMP-1088. The inset shows a pX immunoblot of nanocage proteins released from cells transfected with EPN-pX (Gag myristoylation signal) and VP4-M_7-11_ (HAV VP4 sequence in lieu of the myristoylation signal), with and without IMP-1088 treatment, in an independent experiment. Note that synthesis of the HAV polyprotein can initiate at either the first or second AUG codons (Met) in the ORF.

**Figure S3.**
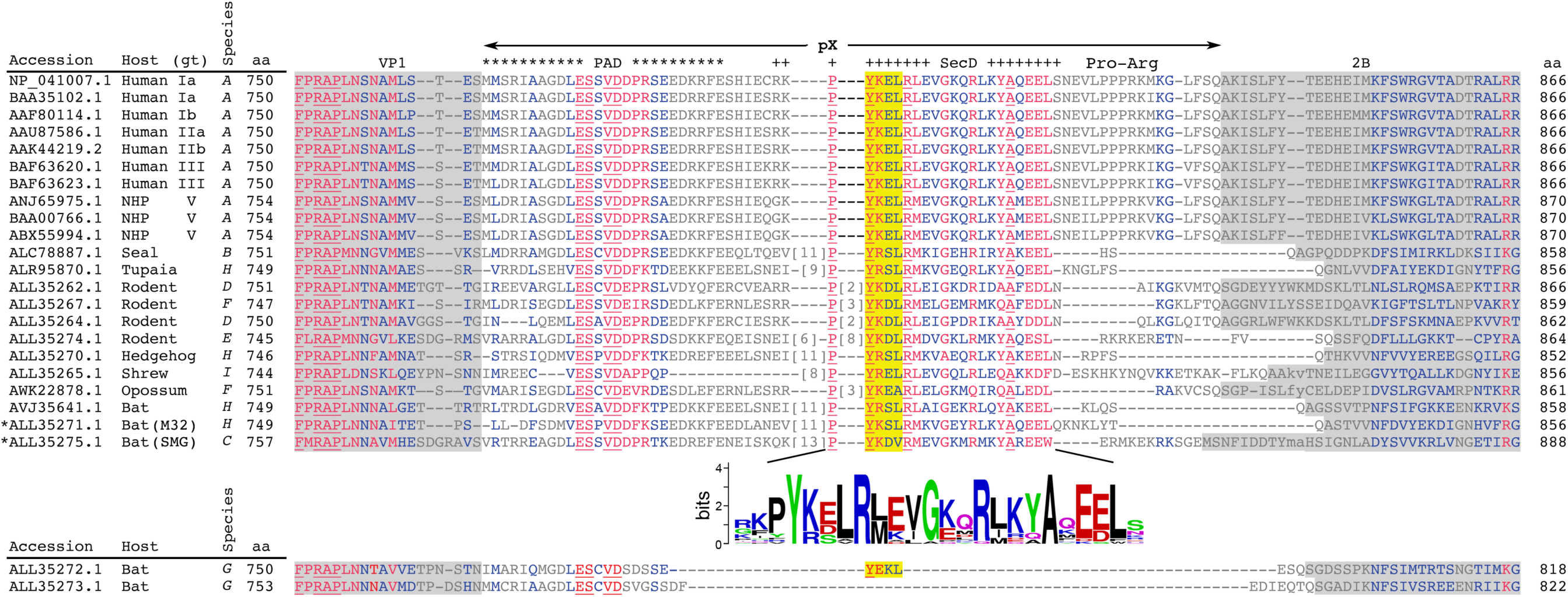
Constraint-based amino acid alignment of pX from 24 near genome-length hepatovirus sequences available in GenBank and recovered from 18 different mammalian species. Residues that are relatively well conserved are shown in red font, those less conserved in blue, followed by very poorly conserved residues in grey font. Residues that are absolutely conserved are underlined. Predicted VP1 and protein 2B sequences flanking pX are shaded in grey; the polyprotein cleavage sites have not been determined experimentally other than for *Hepatovirus A*. Genotype (gt) is noted for human and non-human primate (NHP)-derived strains. The alignment shows relatively high conservation of the secretion domain (SecD, ++++++) which has the motif ‘Y[K/R]xLR[L/M]xxGxxRxxxA’, as well as a second conserved motif (‘[L/V]ESxVD’) within the pentamer assembly domain (PAD, ******). The SecD sequence logo was generated from the 22 sequences above it with WebLogo version 2.8.2 (https://weblogo.berkeley.edu). Two bat *Hepatovirus G* viruses shown at the bottom have a large deletion in the pX region. *Bat virus pX sequences shown to mediate EPN secretion (see Fig. 3B in the main manuscript). The alignment was produced by COBALT, with gap penalties of -11,-1 and a conservation setting of 3 bits (Papadopoulos, J.S., and Agarwala, R. *Bioinformatics* 23: 1073-1079, 2007), then manually adjusted. See **Figure S4A** for a phylogenetic tree.

**Figure S4.**
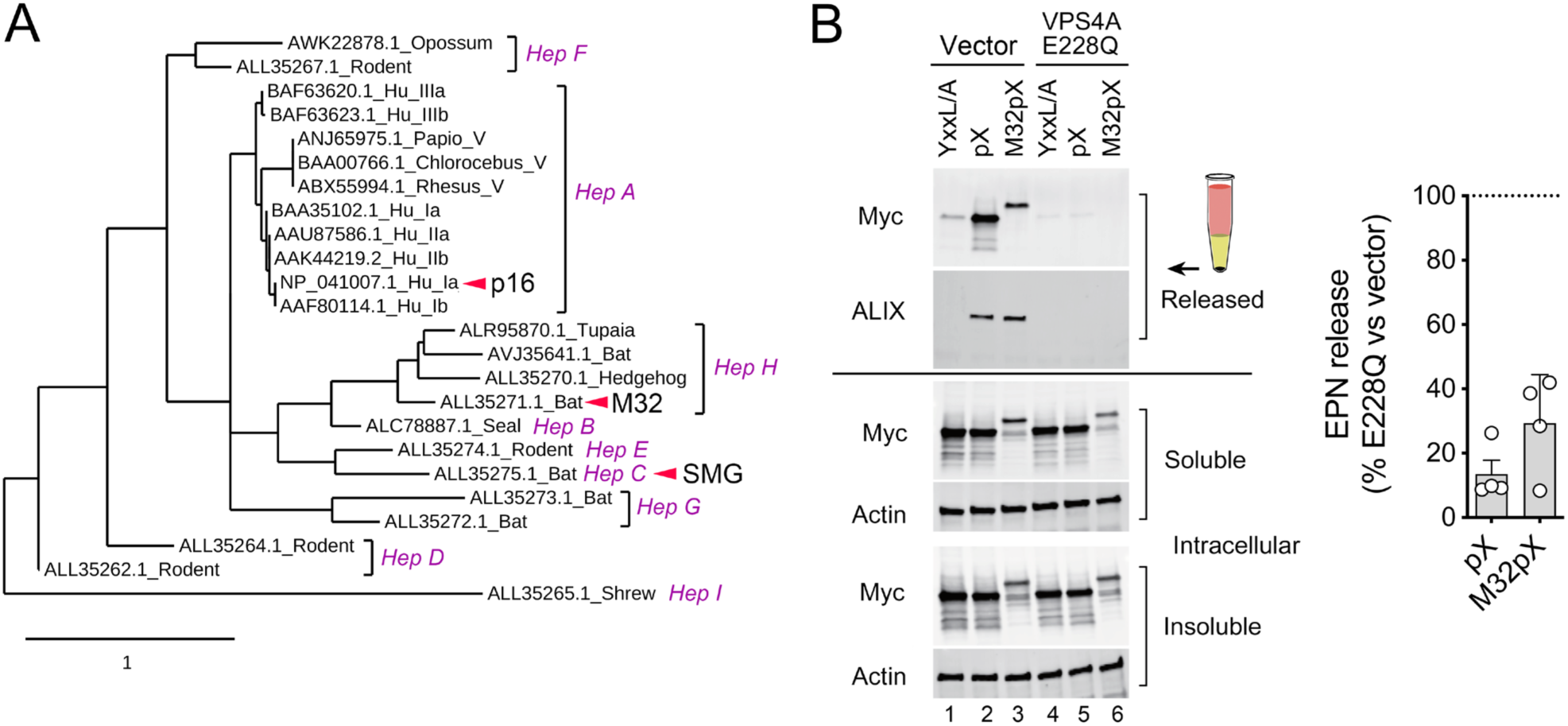
Functional conservation of pX sequences in bat hepatoviruses. **(A)** Phylogenetic tree showing relatedness of amino acid sequences of pX in 9 recognized hepatovirus species infecting 18 different mammalian species (magenta font, “*Hep*” = *Hepatovirus* species). GenBank accession numbers and genotype of *Hepatovirus A* viruses are indicated. Arrows denote pX sequences studied as EPN fusions (Fig. 3 in the main manuscript). (**B**) Immunoblots showing released particulate EPN-pX and EPN-M32pX is associated with ALIX. EPN-YxxL/A was included as a negative control. Cells were co-transfected with an empty vector (lanes 1-3) or vector expressing the dominant-negative VPS4A E228Q mutant (lanes 4-6) to demonstrate that release is ESCRT dependent. To the right is shown the mean normalized percent release of EPN-pX and EPN-M32pX in cells expressing VPSA E228Q versus empty vector, ± S.E.M., n=4.

**Figure S5.**
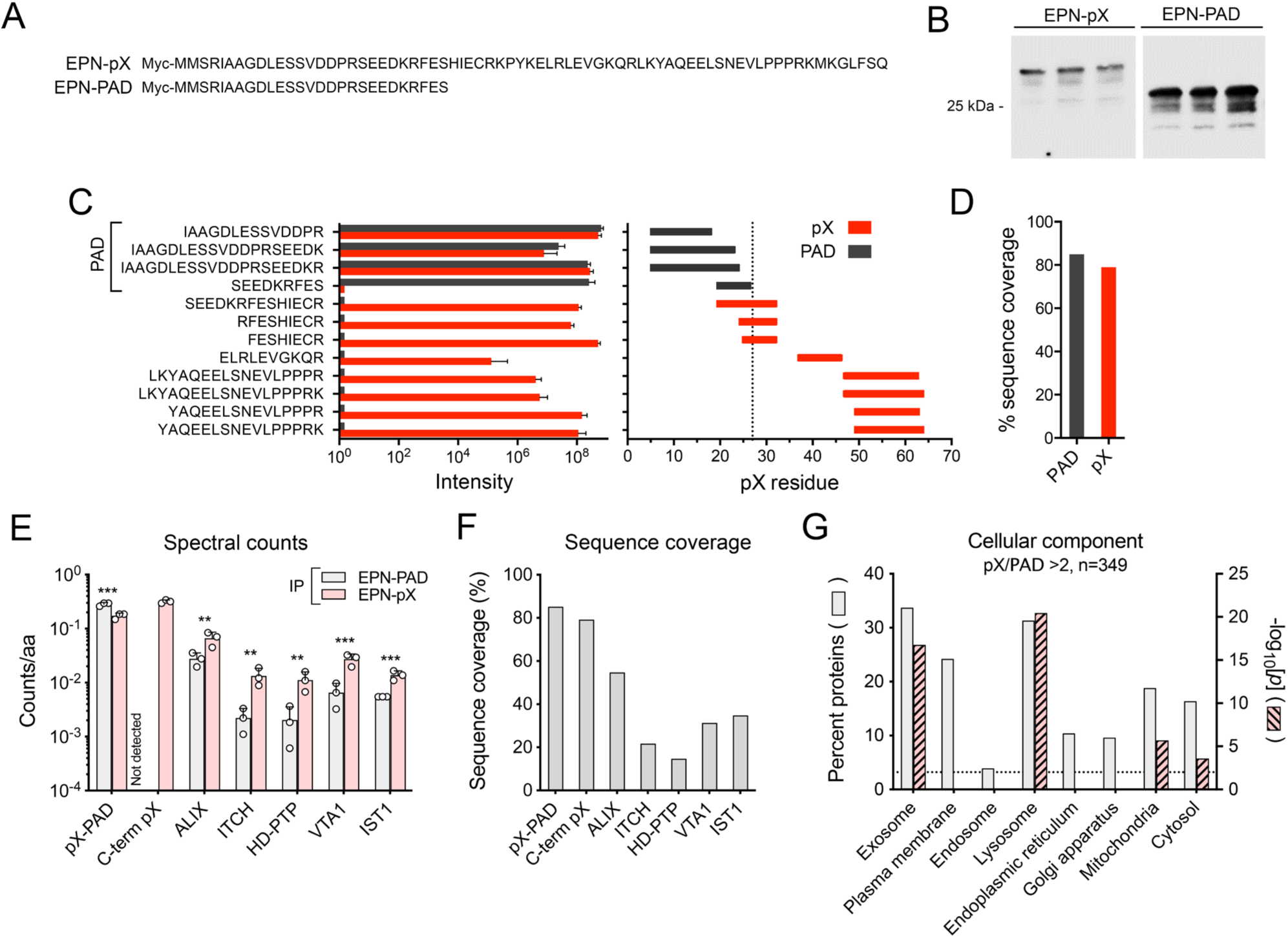
Label-free quantitative (LFQ) proteomic analysis of the C-terminal pX interactome. (**A**) pX sequences in expression vectors used for LFQ analysis of pX-interacting proteins. EPN-PAD contains only the pentamer assembly domain (PAD) of pX. (**B**) Anti-Myc immunoblot showing triplicate protein precipitates subjected to proteomics analysis from 293T cells expressing the EPN-pX and EPN-PAD constructs. (**C**) Mean LFQ intensities of pX peptides in both protein precipitates (n=3 samples of each precipitate, each with 2 technical replicates). Intensities of peptides identified in the EPN-PAD sample (common to both EPN-pX and EPN-PAD) are shown in solid black bars; intensities of peptides unique to EPN-pX are shown in red. (**D**) Percentage of the pX sequence covered by peptides found in the EPN-PAD and EPN-pX samples. (**E**) Spectral counts of selected proteins identified in the proteomics samples. C-term pX = pX sequence uniquely present in EPN-pX and absent in EPN-PAD. **(F)** Peptide coverage of selected proteins identified in the proteomics samples. **(G)** Cellular component of all 349 proteins enriched over 2-fold in EPN-pX compared to EPN-PAD precipitates (FDR<0.05). Significance for protein clustering with each component is plotted on the right axis (hatched bars).

**Figure S6.**
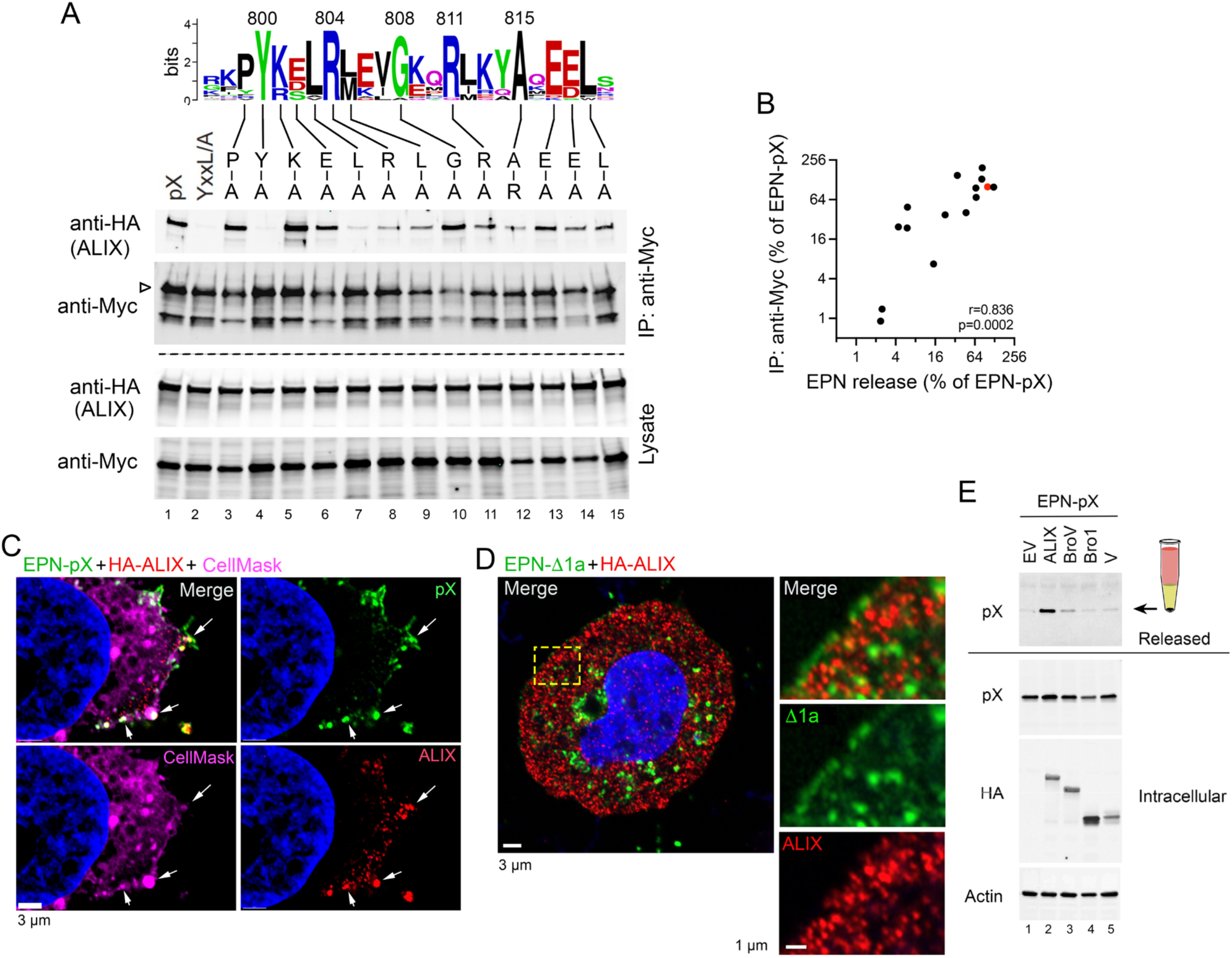
pX interaction with ALIX. **(A)** Co-immunoprecipitation of HA-ALIX and EPN-pX or EPN-pX mutants containing single amino acid substitutions of conserved SecD residues co-expressed in 293T cells. Lysates of 293T cells transfected with DNAs expressing the indicated nanocage proteins and HA-tagged ALIX were precipitated with anti-Myc antibody, then immunoblotted with anti-HA antibody. **(B)** Quantitative comparison of the efficiency of co-immunoprecipitation of EPN constructs with ALIX shown in panel A versus the efficiency of EPN release when fused to pX with mutations in the SecD domain (Fig. 3F). Data are means from 3 (EPN release) or 2 (co-immunoprecipitation) independent experiments, normalized to EPN-pX (100%, red symbol). **(C)** Merged and single-channel Zeiss Airyscan super-resolution fluorescent images of cells transfected with EPN-pX and HA-ALIX expression vectors, with labeling for pX (green) and HA (ALIX, red). Membranes were visualized by labeling with CellMask-647 (magenta). Nuclei were visualized by staining with Hoechst (blue). Extensive pX-ALIX colocalization is evident at buds on the plasma membrane (arrows). **(D)** Super-resolution images of a cell transfected with EPN-Δ1a and HA-ALIX expression vectors showing negligible Δ1a and ALIX colocalization. **(E)** Released extracellular and intracellular EPN-pX nanocage protein in 293T cells overexpressing ALIX or the indicated ALIX domain fragments. EV=empty vector.

**Figure S7.**
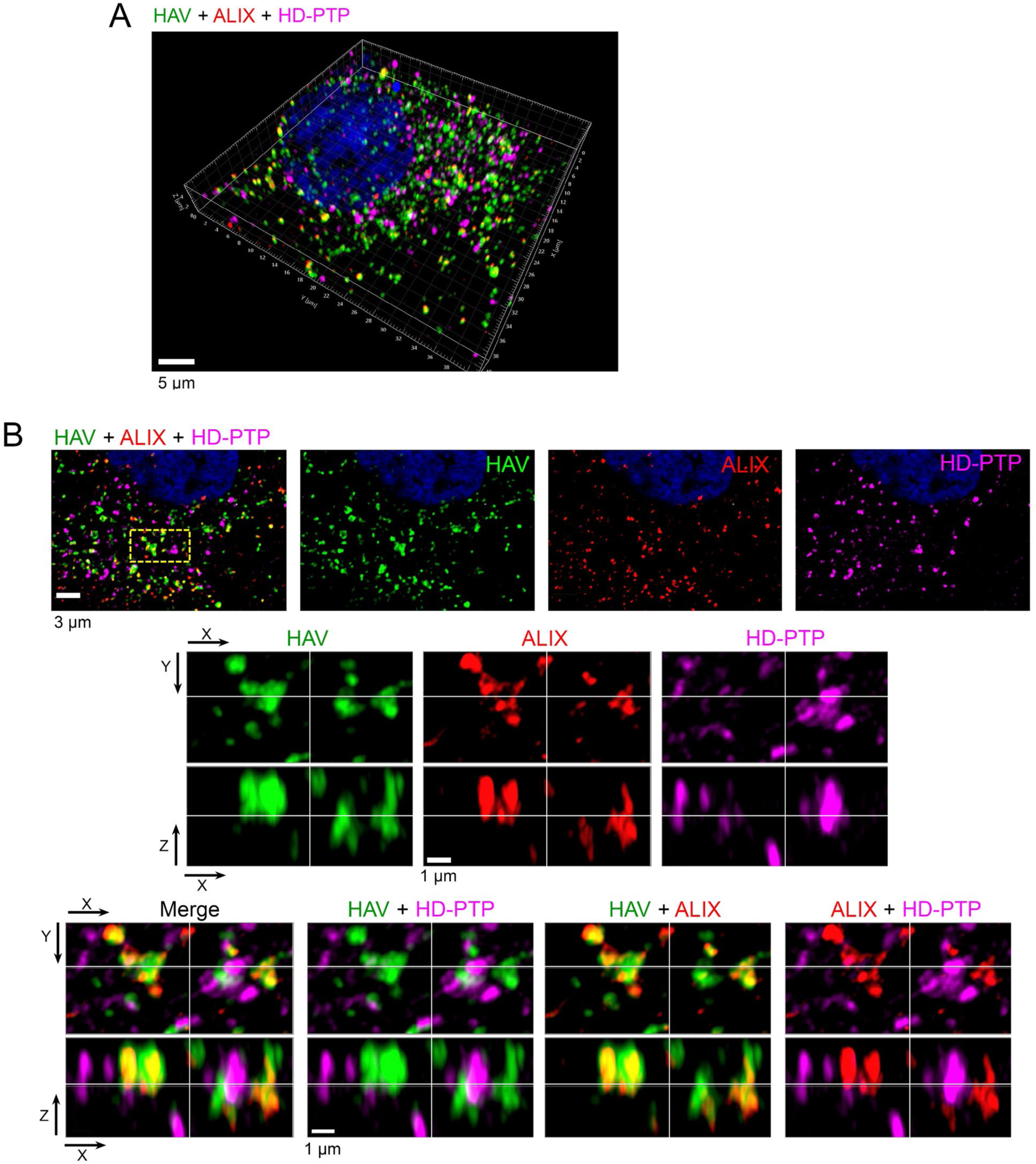
Fluorescence microscopy images of endogenous ALIX and HD-PTP in HAV-infected cells. **(A)** Low magnification view and **(B)** super-resolution fluorescent microscopy images of a cell infected with 18f virus, demonstrating close proximity of viral antigens recognized by polyclonal human anti-HAV (‘HAV’, green), endogenous ALIX (red), and endogenous HD-PTP (magenta). High magnification single- and dual channel images of the region bounded by the dashed yellow box are shown below in three dimensions below.

**Table S1.** Proteins identified by LFQ proteomics as >2-fold enriched in EPN-pX precipitates.

See separate Excel file.

**Table S2.**
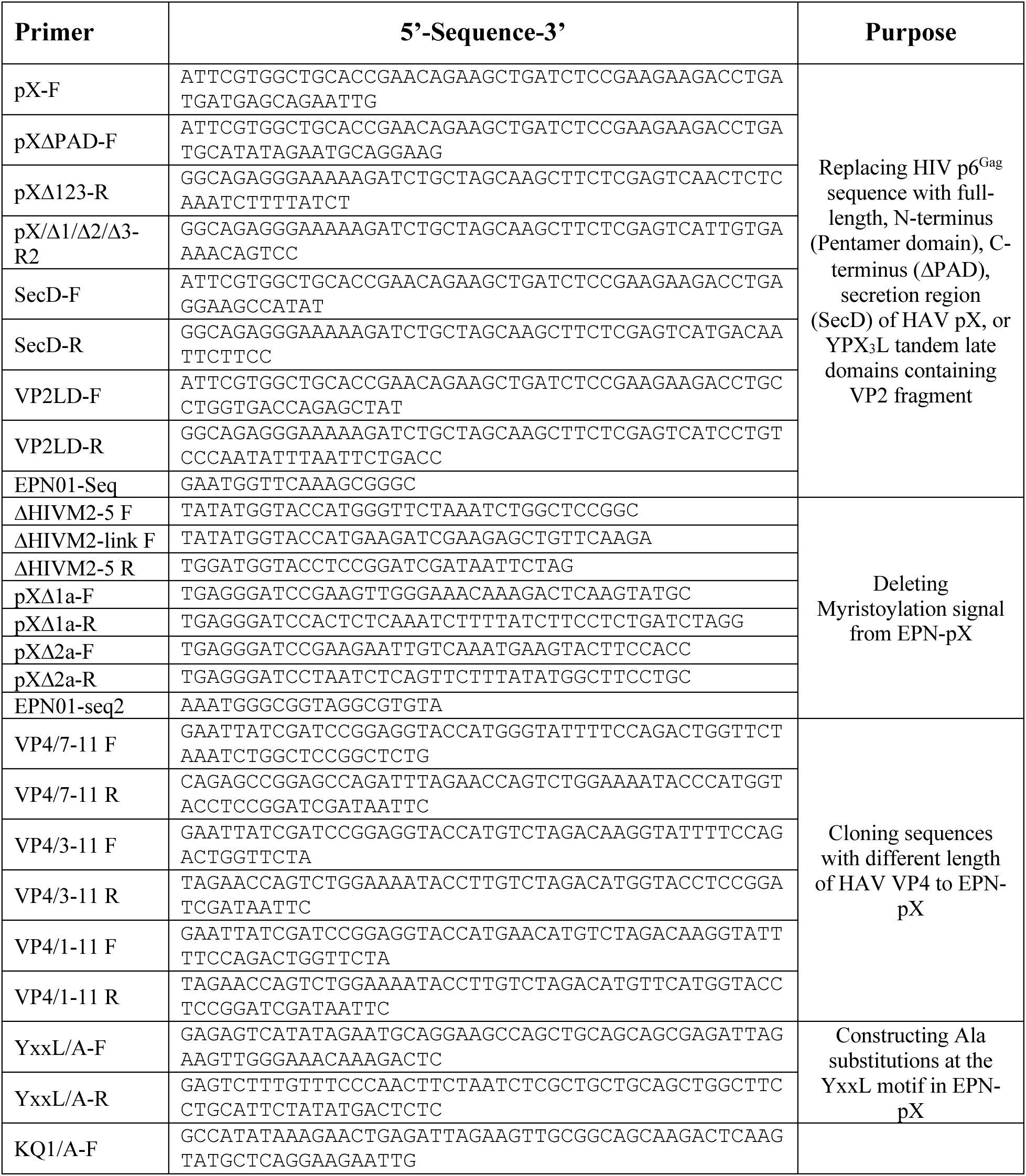
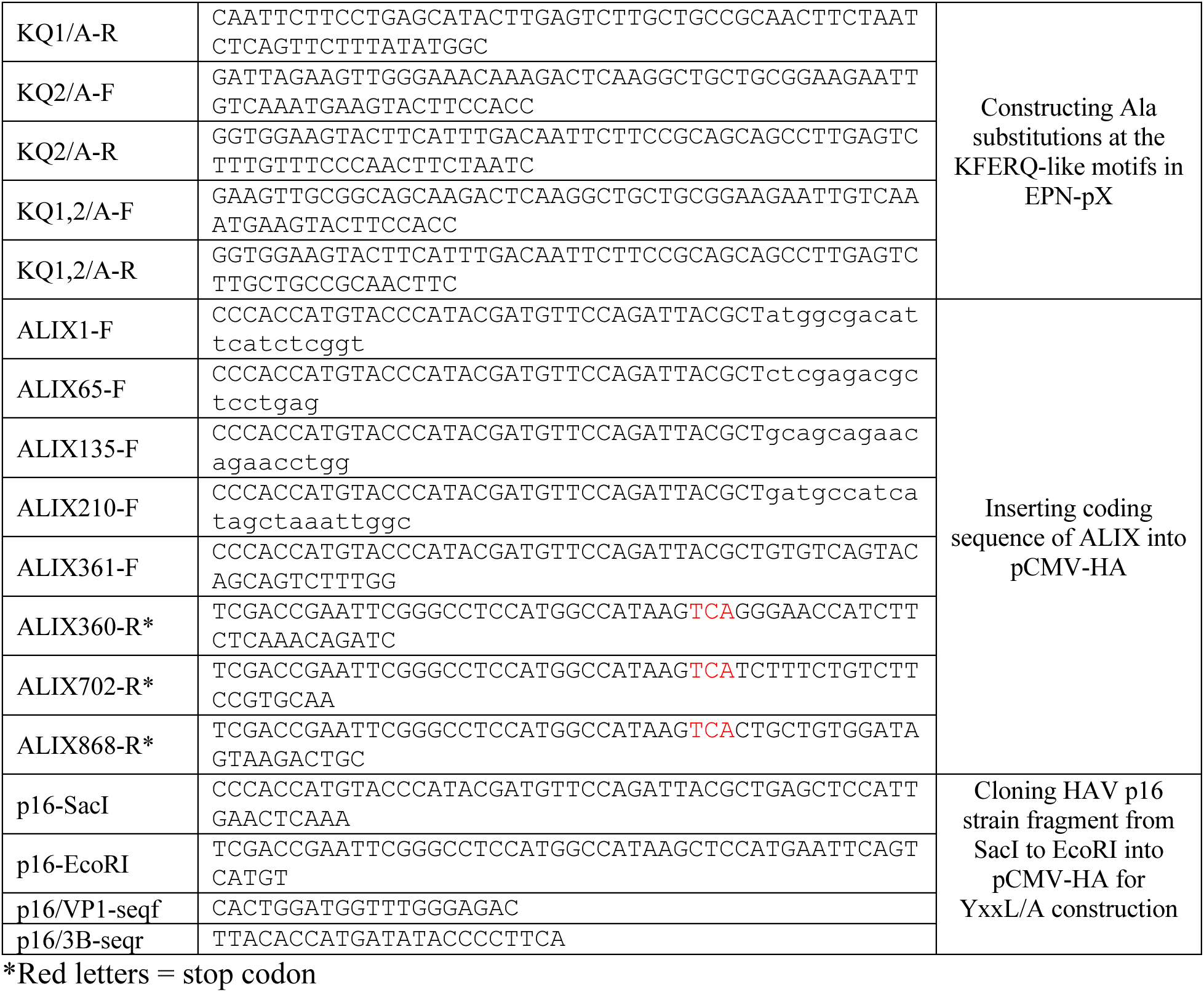
Oligonucleotides used in this study.

**Table S3.**
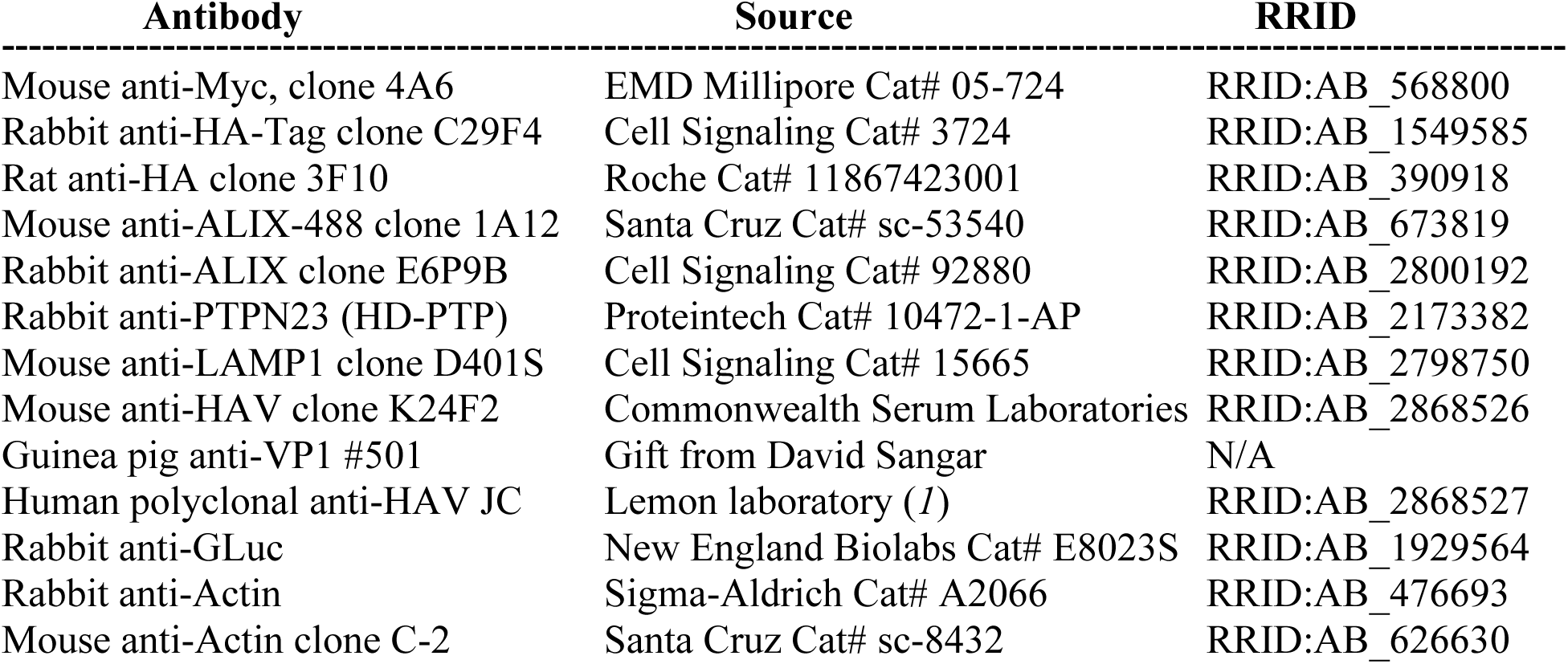
Antibodies used in this study.

## References

1. Feng Z, Hensley L, McKnight KL, Hu F, Madden V, Ping L, et al. A pathogenic picornavirus acquires an envelope by hijacking cellular membranes. Nature. 2013;496(7445):367–71. Epub 2013/04/02. doi: 10.1038/nature12029.

2. Robinson SM, Tsueng G, Sin J, Mangale V, Rahawi S, McIntyre LL, et al. Coxsackievirus B exits the host cell in shed microvesicles displaying autophagosomal markers. PLoS Pathog. 2014;10(4):e1004045. Epub 2014/04/12. doi: 10.1371/journal.ppat.1004045.

3. Santiana M, Ghosh S, Ho BA, Rajasekaran V, Du WL, Mutsafi Y, et al. Vesicle-Cloaked Virus Clusters Are Optimal Units for Inter-organismal Viral Transmission. Cell Host Microbe. 2018;24(2):208–20.e8. Epub 2018/08/10. doi: 10.1016/j.chom.2018.07.006.

4. van der Grein SG, Defourny KAY, Rabouw HH, Galiveti CR, Langereis MA, Wauben MHM, et al. Picornavirus infection induces temporal release of multiple extracellular vesicle subsets that differ in molecular composition and infectious potential. PLoS Pathog. 2019;15(2):e1007594. Epub 2019/02/20. doi: 10.1371/journal.ppat.1007594.

5. Zell R. Picornaviridae-the ever-growing virus family. Arch Virol. 2018;163(2):299–317. Epub 2017/10/24. doi: 10.1007/s00705-017-3614-8.

6. Chen YH, Du W, Hagemeijer MC, Takvorian PM, Pau C, Cali A, et al. Phosphatidylserine vesicles enable efficient en bloc transmission of enteroviruses. Cell. 2015;160(4):619–30. Epub 2015/02/14. doi: 10.1016/j.cell.2015.01.032.

7. McKnight KL, Xie L, Gonzalez-Lopez O, Rivera-Serrano EE, Chen X, Lemon SM. Protein composition of the hepatitis A virus quasi-envelope. Proc Natl Acad Sci U S A. 2017;114(25):6587–92. Epub 2017/05/12. doi: 10.1073/pnas.1619519114.

8. Das A, Hirai-Yuki A, Gonzalez-Lopez O, Rhein B, Moller-Tank S, Brouillette R, et al. TIM1 (HAVCR1) Is not essential for cellular entry of either quasi-enveloped or naked hepatitis A virions. MBio. 2017;8(5): 10.1128/mBio.00969-17. Epub 2017/09/07. doi: 10.1128/mBio.00969-17.

9. Das A, Barrientos RC, Shiota T, Madigan V, Misumi I, McKnight KL, et al. Gangliosides are essential endosomal receptors for quasi-enveloped and naked hepatitis A virus. Nat Microbiol. 2020;5(9):1069–78. doi: doi: 10.1038/s41564-020-0727-8.

10. Hirai-Yuki A, Hensley L, Whitmire JK, Lemon SM. Biliary secretion of quasi-enveloped human hepatitis A virus. MBio. 2016;7:e01998–16.

11. Lemon SM, Ott JJ, Van Damme P, Shouval D. Type A viral hepatitis: A summary and update on the molecular virology, epidemiology, pathogenesis and prevention. J Hepatol. 2018;68:167–84. Epub 2017/09/10. doi: 10.1016/j.jhep.2017.08.034.

12. Hirai-Yuki A, Hensley L, McGivern DR, Gonzalez-Lopez O, Das A, Feng H, et al. MAVS-dependent host species range and pathogenicity of human hepatitis A virus. Science. 2016;353(6307):1541–5. Epub 15 Sep 2016.

13. McCullough J, Frost A, Sundquist WI. Structures, functions, and dynamics of ESCRT-III/Vps4 membrane remodeling and fission complexes. Annu Rev Cell Dev Biol. 2018;34:85–109. Epub 2018/08/11. doi: 10.1146/annurev-cellbio-100616-060600.

14. Vietri M, Radulovic M, Stenmark H. The many functions of ESCRTs. Nat Rev Mol Cell Biol. 2020;21(1):25–42. Epub 2019/11/11. doi: 10.1038/s41580-019-0177-4.

15. Lee S, Joshi A, Nagashima K, Freed EO, Hurley JH. Structural basis for viral late-domain binding to Alix. Nat Struct Mol Biol. 2007;14(3):194–9. Epub 2007/02/06. doi: 10.1038/nsmb1203.

16. Gonzalez-Lopez O, Rivera-Serrano EE, Hu F, Hensley L, McKnight KL, Ren J, et al. Redundant late domain functions of tandem VP2 YPX3L motifs in nonlytic cellular egress of quasi-enveloped hepatitis A virus. J Virol. 2018;92(23). Epub 2018/09/21. doi: 10.1128/JVI.01308-18.

17. Votteler J, Sundquist WI. Virus budding and the ESCRT pathway. Cell Host Microbe. 2013;14(3):232–41. Epub 2013/09/17. doi: 10.1016/j.chom.2013.08.012.

18. Ren X, Hurley JH. Proline-rich regions and motifs in trafficking: from ESCRT interaction to viral exploitation. Traffic. 2011;12(10):1282–90. Epub 2011/04/27. doi: 10.1111/j.1600-0854.2011.01208.x [doi].

19. Wang X, Ren J, Gao Q, Hu Z, Sun Y, Li X, et al. Hepatitis A virus and the origins of picornaviruses. Nature. 2015;517(7532):85–8. Epub 2014/10/21. doi: 10.1038/nature13806.

20. Anderson DA, Ross BC. Morphogenesis of hepatitis A virus: Isolation and characterization of subviral particles. J Virol. 1990;64:5284–9.

21. Cohen L, Benichou D, Martin A. Analysis of deletion mutants indicates that the 2A polypeptide of hepatitis A virus participates in virion morphogenesis. J Virol. 2002;76(15):7495–505.

22. Probst C, Jecht M, Gauss-Muller V. Intrinsic signals for the assembly of hepatitis A virus particles. Role of structural proteins VP4 and 2A. J Biol Chem. 1999;274(8):4527–31.

23. Harmon SA, Emerson SU, Huang YK, Summers DF, Ehrenfeld E. Hepatitis A viruses with deletions in the 2A gene are infectious in cultured cells and marmosets. J Virol. 1995;69(Sep):5576–81.

24. Jiang W, Ma P, Deng L, Liu Z, Wang X, Liu X, et al. Hepatitis A virus structural protein pX interacts with ALIX and promotes the secretion of virions and foreign proteins through exosome-like vesicles. J Extracell Vesicles. 2020;9(1):1716513. Epub 2020/02/23. doi: 10.1080/20013078.2020.1716513.

25. Desrochers G, Kazan JM, Pause A. Structure and functions of His domain protein tyrosine phosphatase in receptor trafficking and cancer (1). Biochem Cell Biol. 2019;97(1):68–72. Epub 2018/06/08. doi: 10.1139/bcb-2017-0322.

26. Neveu G, Barouch-Bentov R, Ziv-Av A, Gerber D, Jacob Y, Einav S. Identification and targeting of an interaction between a tyrosine motif within hepatitis C virus core protein and AP2M1 essential for viral assembly. PLoS Pathog. 2012;8(8):e1002845. Epub 2012/08/24. doi: 10.1371/journal.ppat.1002845.

27. Barouch-Bentov R, Neveu G, Xiao F, Beer M, Bekerman E, Schor S, et al. Hepatitis C virus proteins interact with the endosomal sorting complex required for transport (ESCRT) machinery via ubiquitination to facilitate viral envelopment. MBio. 2016;7(6):10.1128/mBio.01456-16. Epub 2016/11/03. doi: 10.1128/mBio.01456-16.

28. Cassonnet P, Rolloy C, Neveu G, Vidalain PO, Chantier T, Pellet J, et al. Benchmarking a luciferase complementation assay for detecting protein complexes. Nat Methods. 2011;8(12):990–2. Epub 2011/12/01. doi: 10.1038/nmeth.1773.

29. Hsia Y, Bale JB, Gonen S, Shi D, Sheffler W, Fong KK, et al. Design of a hyperstable 60-subunit protein dodecahedron. [corrected]. Nature. 2016;535(7610):136–9. Epub 2016/06/17. doi: 10.1038/nature18010.

30. Votteler J, Ogohara C, Yi S, Hsia Y, Nattermann U, Belnap DM, et al. Designed proteins induce the formation of nanocage-containing extracellular vesicles. Nature. 2016;540(7632):292–5. Epub 2016/12/06. doi: 10.1038/nature20607.

31. Panjwani A, Strauss M, Gold S, Wenham H, Jackson T, Chou JJ, et al. Capsid protein VP4 of human rhinovirus induces membrane permeability by the formation of a size-selective multimeric pore. PLoS Pathog. 2014;10(8):e1004294. Epub 2014/08/08. doi: 10.1371/journal.ppat.1004294.

32. Shukla A, Padhi AK, Gomes J, Banerjee M. The VP4 peptide of hepatitis A virus ruptures membranes through formation of discrete pores. J Virol. 2014;88(21):12409–21. Epub 2014/08/15. doi: 10.1128/jvi.01896-14.

33. Mousnier A, Bell AS, Swieboda DP, Morales-Sanfrutos J, Perez-Dorado I, Brannigan JA, et al. Fragment-derived inhibitors of human N-myristoyltransferase block capsid assembly and replication of the common cold virus. Nat Chem. 2018;10(6):599–606. Epub 2018/05/16. doi: 10.1038/s41557-018-0039-2.

34. Wang X, Ren J, Gao Q, Hu Z, Sun Y, Li X, et al. Hepatitis A virus and the origins of picornaviruses. Nature. 2015;517(7532):85–8. Epub 2014 Oct 19.

35. Lin J, Lee LY, Roivainen M, Filman DJ, Hogle JM, Belnap DM. Structure of the Fab-labeled “breathing” state of native poliovirus. J Virol. 2012;86(10):5959–62. Epub 2012/03/09. doi: 10.1128/jvi.05990-11.

36. Shtanko O, Watanabe S, Jasenosky LD, Watanabe T, Kawaoka Y. ALIX/AIP1 is required for NP incorporation into Mopeia virus Z-induced virus-like particles. J Virol. 2011;85(7):3631–41. Epub 2011/01/21. doi: 10.1128/JVI.01984-10.

37. Wolff S, Ebihara H, Groseth A. Arenavirus budding: a common pathway with mechanistic differences. Viruses. 2013;5(2):528–49. Epub 2013/02/26. doi: 10.3390/v5020528.

38. Irie T, Shimazu Y, Yoshida T, Sakaguchi T. The YLDL sequence within Sendai virus M protein is critical for budding of virus-like particles and interacts with Alix/AIP1 independently of C protein. J Virol. 2007;81(5):2263–73. Epub 2006/12/15. doi: 10.1128/JVI.02218-06.

39. Smith DB, Simmonds P. Classification and Genomic Diversity of Enterically Transmitted Hepatitis Viruses. Cold Spring Harb Perspect Med. 2018;8(9):a031880. Epub 2018/03/14. doi: 10.1101/cshperspect.a031880.

40. Drexler JF, Corman VM, Lukashev AN, van den Brand JMA, Gmyl A, Brunink S, et al. Evolutionary origins of hepatitis A virus in small mammals. Proc Nat’l Acad Sci U S A. 2015;112:15190–5. Epub 2015 Nov 2.

41. Anthony SJ, St Leger JA, Liang E, Hicks AL, Sanchez-Leon MD, Jain K, et al. Discovery of a novel Hepatovirus (phopivirus of seals) related to human hepatitis A virus. MBio. 2015;6(4):10.1128/mBio.01180-15. Epub 2015/08/27. doi: 10.1128/mBio.01180-15.

42. Tabernero L, Woodman P. Dissecting the role of His domain protein tyrosine phosphatase/PTPN23 and ESCRTs in sorting activated epidermal growth factor receptor to the multivesicular body. Biochem Soc Trans. 2018;46(5):1037–46. Epub 2018/09/08. doi: 10.1042/bst20170443.

43. Lee J, Oh KJ, Lee D, Kim BY, Choi JS, Ku B, et al. Structural study of the HD-PTP Bro1 domain in a complex with the core region of STAM2, a subunit of ESCRT-0. PLoS One. 2016;11(2):e0149113. Epub 2016/02/13. doi: 10.1371/journal.pone.0149113.

44. Doyotte A, Mironov A, McKenzie E, Woodman P. The Bro1-related protein HD-PTP/PTPN23 is required for endosomal cargo sorting and multivesicular body morphogenesis. Proc Natl Acad Sci U S A. 2008;105(17):6308–13. Epub 2008/04/25. doi: 10.1073/pnas.0707601105.

45. Xiao J, Xia H, Zhou J, Azmi IF, Davies BA, Katzmann DJ, et al. Structural basis of Vta1 function in the multivesicular body sorting pathway. Dev Cell. 2008;14(1):37–49. Epub 2008/01/16. doi: 10.1016/j.devcel.2007.10.013.

46. Ziegler CM, Dang L, Eisenhauer P, Kelly JA, King BR, Klaus JP, et al. NEDD4 family ubiquitin ligases associate with LCMV Z’s PPXY domain and are required for virus budding, but not via direct ubiquitination of Z. PLoS Pathog. 2019;15(11):e1008100. Epub 2019/11/12. doi: 10.1371/journal.ppat.1008100.

47. Han Z, Sagum CA, Bedford MT, Sidhu SS, Sudol M, Harty RN. ITCH E3 Ubiquitin Ligase Interacts with Ebola Virus VP40 To Regulate Budding. J Virol. 2016;90(20):9163–71. Epub 2016/08/05. doi: 10.1128/jvi.01078-16.

48. Fisher RD, Chung HY, Zhai Q, Robinson H, Sundquist WI, Hill CP. Structural and biochemical studies of ALIX/AIP1 and its role in retrovirus budding. Cell. 2007;128(5):841–52. Epub 2007/03/14. doi: 10.1016/j.cell.2007.01.035.

49. Kim J, Sitaraman S, Hierro A, Beach BM, Odorizzi G, Hurley JH. Structural basis for endosomal targeting by the Bro1 domain. Dev Cell. 2005;8(6):937–47. Epub 2005/06/07. doi: 10.1016/j.devcel.2005.04.001.

50. Zhai Q, Landesman MB, Robinson H, Sundquist WI, Hill CP. Structure of the Bro1 domain protein BROX and functional analyses of the ALIX Bro1 domain in HIV-1 budding. PLoS ONE. 2011;6(12):e27466. Epub 2011/12/14. doi: 10.1371/journal.pone.0027466.

51. McCullough J, Fisher RD, Whitby FG, Sundquist WI, Hill CP. ALIX-CHMP4 interactions in the human ESCRT pathway. Proc Natl Acad Sci U S A. 2008;105(22):7687–91. Epub 2008/05/31. doi: 10.1073/pnas.0801567105.

52. Gahloth D, Heaven G, Jowitt TA, Mould AP, Bella J, Baldock C, et al. The open architecture of HD-PTP phosphatase provides new insights into the mechanism of regulation of ESCRT function. Sci Rep. 2017;7(1):9151. Epub 2017/08/24. doi: 10.1038/s41598-017-09467-9.

53. Vives-Adrian L, Garriga D, Buxaderas M, Fraga J, Pereira PJ, Macedo-Ribeiro S, et al. Structural basis for host membrane remodeling induced by protein 2B of hepatitis A virus. J Virol. 2015;89(7):3648–58. Epub 2015/01/16. doi: 10.1128/jvi.02881-14.

54. Bissig C, Gruenberg J. ALIX and the multivesicular endosome: ALIX in Wonderland. Trends Cell Biol. 2014;24(1):19–25. Epub 2013/11/30. doi: S0962-8924(13)00190-6 [pii] 10.1016/j.tcb.2013.10.009 [doi].

55. Skowyra ML, Schlesinger PH, Naismith TV, Hanson PI. Triggered recruitment of ESCRT machinery promotes endolysosomal repair. Science. 2018;360(6384). Epub 2018/04/07. doi: 10.1126/science.aar5078.

56. Parkinson G, Roboti P, Zhang L, Taylor S, Woodman P. His domain protein tyrosine phosphatase and Rabaptin-5 couple endo-lysosomal sorting of EGFR with endosomal maturation. J Cell Sci. 2021;134(21). Epub 20211104. doi: 10.1242/jcs.259192.

57. Popov S, Popova E, Inoue M, Göttlinger HG. Divergent Bro1 domains share the capacity to bind human immunodeficiency virus type 1 nucleocapsid and to enhance virus-like particle production. J Virol. 2009;83(14):7185–93. Epub 2009/05/01. doi: 10.1128/jvi.00198-09.

58. Zhai Q, Fisher RD, Chung HY, Myszka DG, Sundquist WI, Hill CP. Structural and functional studies of ALIX interactions with YPX(n)L late domains of HIV-1 and EIAV. Nat Struct Mol Biol. 2008;15(1):43–9. Epub 2007/12/11. doi: 10.1038/nsmb1319.

59. Zhang S, Fan G, Hao Y, Hammell M, Wilkinson JE, Tonks NK. Suppression of protein tyrosine phosphatase N23 predisposes to breast tumorigenesis via activation of FYN kinase. Genes Dev. 2017;31(19):1939–57. Epub 20171024. doi: 10.1101/gad.304261.117.

60. Oda K, Matoba Y, Sugiyama M, Sakaguchi T. Structural Insight into the Interaction of Sendai Virus C Protein with Alix To Stimulate Viral Budding. J Virol. 2021;95(19):e0081521. Epub 20210909. doi: 10.1128/jvi.00815-21.

61. Chung HY, Morita E, von Schwedler U, Müller B, Kräusslich HG, Sundquist WI. NEDD4L overexpression rescues the release and infectivity of human immunodeficiency virus type 1 constructs lacking PTAP and YPXL late domains. J Virol. 2008;82(10):4884–97. Epub 2008/03/07. doi: 10.1128/jvi.02667-07.

62. Baillet N, Krieger S, Carnec X, Mateo M, Journeaux A, Merabet O, et al. E3 Ligase ITCH Interacts with the Z Matrix Protein of Lassa and Mopeia Viruses and Is Required for the Release of Infectious Particles. Viruses. 2019;12(1):49. Epub 2020/01/08. doi: 10.3390/v12010049.

63. Keren-Kaplan T, Attali I, Estrin M, Kuo LS, Farkash E, Jerabek-Willemsen M, et al. Structure-based in silico identification of ubiquitin-binding domains provides insights into the ALIX-V:ubiquitin complex and retrovirus budding. EMBO J. 2013;32(4):538–51. Epub 20130129. doi: 10.1038/emboj.2013.4.

64. Pashkova N, Yu L, Schnicker NJ, Tseng CC, Gakhar L, Katzmann DJ, et al. Interactions of ubiquitin and CHMP5 with the V domain of HD-PTP reveals role for regulation of Vps4 ATPase. Mol Biol Cell. 2021;32(22):ar42. Epub 20210929. doi: 10.1091/mbc.E21-04-0219.

65. Blight KJ, McKeating JA, Rice CM. Highly permissive cell lines for subgenomic and genomic hepatitis C virus RNA replication. J Virol. 2002;76(24):13001–14.

66. Feng H, Lenarcic EM, Yamane D, Wauthier E, Mo J, Guo H, et al. NLRX1 promotes immediate IRF1-directed antiviral responses by limiting dsRNA-activated translational inhibition mediated by PKR. Nat Immunol. 2017;18(12):1299–309. Epub 2017/10/03. doi: 10.1038/ni.3853.

67. Jansen RW, Newbold JE, Lemon SM. Complete nucleotide sequence of a cell culture-adapted variant of hepatitis A virus: comparison with wild-type virus with restricted capacity for in vitro replication. Virology. 1988;163:299–307.

68. Lemon SM, Murphy PC, Shields PA, Ping LH, Feinstone SM, Cromeans T, et al. Antigenic and genetic variation in cytopathic hepatitis A virus variants arising during persistent infection: evidence for genetic recombination. J Virol. 1991;65:2056–65.

69. Yamane D, Feng H, Rivera-Serrano EE, Selitsky SR, Hirai-Yuki A, Das A, et al. Constitutive expression of interferon regulatory factor 1 drives intrinsic hepatocyte resistance to multiple RNA viruses. Nat Microbiol. 2019;4(7):1096–104.

70. Yamane D, McGivern DR, Wauthier E, Yi M, Madden VJ, Welsch C, et al. Regulation of the hepatitis C virus RNA replicase by endogenous lipid peroxidation. Nat Med. 2014;20(8):927–35. Epub 2014/07/30. doi: 10.1038/nm.3610.

71. Bachurski D, Schuldner M, Nguyen PH, Malz A, Reiners KS, Grenzi PC, et al. Extracellular vesicle measurements with nanoparticle tracking analysis - An accuracy and repeatability comparison between NanoSight NS300 and ZetaView. J Extracell Vesicles. 2019;8(1):1596016. Epub 2019/04/17. doi: 10.1080/20013078.2019.1596016.

72. Boussif O, Lezoualc’h F, Zanta MA, Mergny MD, Scherman D, Demeneix B, et al. A versatile vector for gene and oligonucleotide transfer into cells in culture and in vivo: polyethylenimine. Proc Natl Acad Sci U S A. 1995;92(16):7297–301. Epub 1995/08/01. doi: 10.1073/pnas.92.16.7297.

73. Zheng SQ, Palovcak E, Armache JP, Verba KA, Cheng Y, Agard DA. MotionCor2: anisotropic correction of beam-induced motion for improved cryo-electron microscopy. Nat Methods. 2017;14(4):331–2. Epub 2017/03/03. doi: 10.1038/nmeth.4193.

74. Rohou A, Grigorieff N. CTFFIND4: Fast and accurate defocus estimation from electron micrographs. J Struct Biol. 2015;192(2):216–21. Epub 2015/08/19. doi: 10.1016/j.jsb.2015.08.008.

75. Zivanov J, Nakane T, Forsberg BO, Kimanius D, Hagen WJ, Lindahl E, et al. New tools for automated high-resolution cryo-EM structure determination in RELION-3. Elife. 2018;7:10.7554/eLife.42166. Epub 2018/11/10. doi: 10.7554/eLife.42166.

